# Anchoring of parasitic plasmids to inactive regions of eukaryotic chromosomes through nucleosome signal

**DOI:** 10.1101/2023.10.04.558402

**Authors:** Fabien Girard, Antoine Even, Agnès Thierry, Myriam Ruault, Léa Meneu, Sandrine Adiba, Angela Taddei, Romain Koszul, Axel Cournac

## Abstract

Natural plasmids are common in prokaryotes but few have been documented in eukaryotes. The natural 2µ plasmid present in budding yeast *Saccharomyces cerevisiae* is one of the most well characterized. This highly stable genetic element coexists with its host for millions of years, efficiently segregating at each cell division through a mechanism that remains poorly understood. Using proximity ligation (Hi-C, Micro-C) to map the contacts between the 2µ and yeast chromosomes under dozens of different biological conditions, we found that the plasmid tether preferentially on regions with low transcriptional activity, often corresponding to long inactive genes. Common players in chromosome structure such as members of the structural maintenance of chromosome complexes (SMC) are not involved in these contacts which depend instead on a nucleosomal signal associated with a depletion of RNA Pol II. These contacts are stable throughout the cell cycle, and can be established within minutes. This strategy may involve other types of DNA molecules and species other than *S. cerevisiae*, as suggested by the binding pattern of the natural plasmid along the silent regions of the chromosomes of *Dictyostelium discoideum*.

## Introduction

The way in which mobile DNA elements, such as plasmids, viruses and transposons, are maintained within their hosts is a key question for understanding their phenotypes and their impact. Many eukaryotic DNA viruses are pathogens that can represent public health issues Some can even be retained over a long period of time in the nucleus through several mechanisms. For instance, the hepatitis B virus (HBV) can integrate into the genome and remain in a latent state for very long periods (Dias et al., 2022). The Epstein Barr virus (EBV) of the herpesvirus family remains as an episome in the nucleus able to replicate and hitchhike on host chromosomes thus vertically transferred throughout cell division (Coursey and McBride, 2019; Kim et al., 2020). In contrast, few examples of plasmids naturally present in eukaryotic nuclei have been documented (Esser et al., 2012). To our knowledge, only two families of natural plasmid present in eukaryotic nuclei have been clearly identified and characterized: the *Saccharomyces cerevisiae* 2μ plasmid (Sau et al., 2019) and derivative in other ascomycota species, and the Ddp plasmid family in the social amoeba *Dictyostelium discoideum* (Rieben et al., 1998; Shammat et al., 1998). Why such a small number of plasmids remains unclear: either these objects have been less studied and are overlooked in sequencing data, or eukaryotes have developed sufficiently effective protection and/or defense mechanisms. In either case, it is not yet clear how these molecules are maintained across generations.

The 2μ plasmid is one of the most studied examples of a selfish DNA element, i.e. a molecule that does not appear to confer any fitness advantage on its host, without imposing a significant cost (Mead et al., 1986). The 6.3 kb sequence, named after its contour length when observed with electron microscopy (Sinclair et al. 1967), is replicated by the host replication machinery (Zakian et al., 1979). It is present in most *S. cerevisiae* natural isolates and laboratory strains (Peter et al., 2018) suggesting a very efficient and successful persistence mechanism. It encodes a partitioning system that includes Rep1 and Rep2, two proteins that associate with the plasmid STB repeat sequence that is essential for plasmid stability (McQuaid et al., 2019) and a specific recombinase Flp1. The origin of this plasmid is not well known. Various clues could point to the origin of bacteriophages. The specific Flp1 recombinase is a tyrosine recombinase like the Cre recombinase of bacteriophage P1. It has also been shown that bacteria can naturally transform *S. cerevisiae* yeast by conjugation (Heinemann and Sprague, 1989) making possible a transfer between different species of genetic material that evolved into the 2µ plasmid we know today.

Several works point at a ‘chromosome hitchhiking’ mechanism by which the plasmid binds to the chromosomes of the host, taking advantage of its segregation machinery during cell divisions (Sau et al., 2019). Microscopy investigations show that the 2µ plasmid colocalizes with the host chromatin, though its precise nuclear localisation remains unclear, with studies suggesting either a preferential position at the center of the nucleus (Heun et al., 2001), or in nuclear periphery close to telomeres (Kumar et al., 2021).

De novo calling of DNA contacts between molecules is difficult to achieve with microscopy but can be done using capture of chromosome conformation approaches such as Hi-C (Lieberman-Aiden et al., 2009, Hsieh et al., 2016). We therefore used Hi-C and Micro-C data to map and quantify the plasmid physical contacts with the host chromosomes. We show that the plasmid contacts a set of discrete loci that remain remarkably stable in multiple growth and mutant conditions. The set of contact hotspots corresponds to relatively long, inactive regions. These contacts can evolve within a few minutes following environmental stress, and depend on the nucleosomal H4 basic patch. Strikingly, a Mb long inactive artificial bacterial chromosome was able to bind plasmid molecules from their native binding positions, illustrating how the plasmid is spontaneously attracted by inactive chromatin, whatever its origin. Overall, these results point to the existence of a segregation mechanism whereby the plasmid may recognize a signal associated with chromatin structure to bind to the inactive chromatin regions of its hosts in a reversible way.

Very few eukaryotic plasmids have been identified. Other Saccharomycetaceae, such as the Lachancea species *Lachancea fermentati* and *waltii* that have diverged from Saccharomyces for over 100 My ago also display episomes homologous to the 2µ. Further in the phylogenetic tree, the amoeba *Dictyostelium discoideum* contains the Ddp5 plasmid. We show that these episomes also preferentially tether to inactive regions, suggesting that this may be a widespread strategy for eukaryotic plasmids to ensure their correct segregation during cell division in a way that does not disrupt host regulation.

## Results

### Chromatinization and 3D folding of the 2µ plasmid

The 2µ plasmid sequence is typically filtered (or overlooked) from Hi-C analysis and more generally from any high-throughput sequencing data. We therefore revisited ten years of datasets to explore this overlooked sequence. Using Micro-C, a high-resolution Hi-C derivative that quantifies DNA contacts at the nucleosomal level (Hsieh et al., 2016; Swygert et al., 2019), we generated a plasmid contact map at 200 bp resolution from cells in exponential growth (**Fig. 1a**; **Methods**). The cis-contacts map revealed four small self-interacting regions corresponding to the four plasmid genes, reminiscent of the pattern observed at the level of active chromosomal genes (Hsieh et al., 2016). The ∼1 kb STB-ORI region, positioned in between the RAF1 and REP2 genes, appeared as a constrained region with a stronger local enrichment in short range contacts.

**Figure 1.**
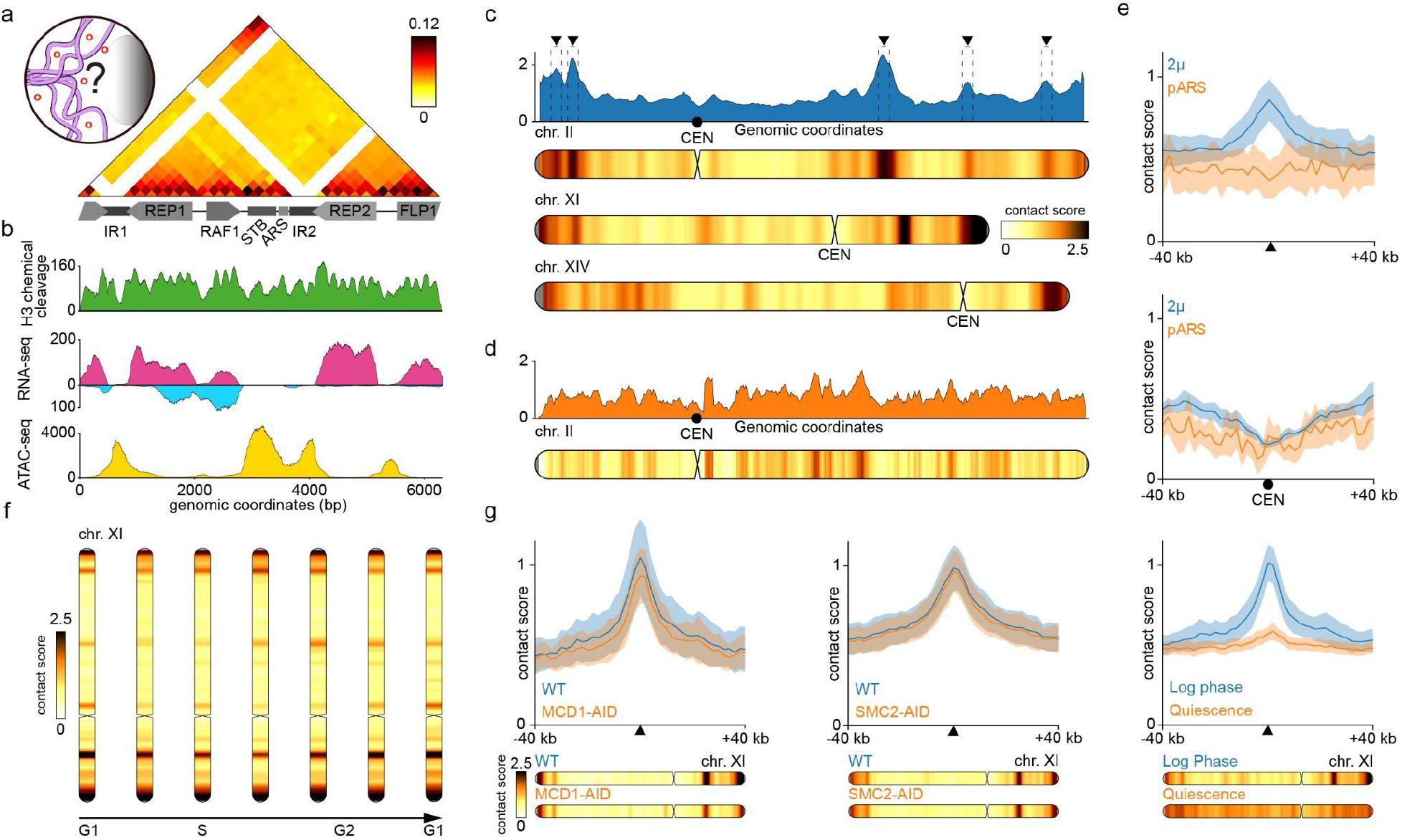
Specific positioning of the natural 2µ plasmid on several genomic regions of *S. cerevisiae* chromosomes. **a,** Contact map of the 2µ plasmid with bins of 200 bp based on Micro-C XL data in asynchronous cells (log phase). The 4 genes and cis-acting sequences of the 2µ plasmid genome are annotated below the map **b,** MNase-seq giving the nucleosome density, RNA-seq giving transcription level on forward and reverse strands and ATAC-seq giving chromatin accessibility along the 2µ plasmid. **c,** Contact profile of 2µ plasmid with several chromosomes of *S. cerevisiae* (binned at 2 kb). **d,** Contact profile of pARS plasmid (containing no 2μ or centromere systems) for chromosome II. **e,** Averaged contact signal of the 2µ and pARS plasmids at the positions automatically detected in WT, log phase for 2µ plasmid (top) and at centromeres positions (below). **f,** Contact profile of the 2µ plasmid with chr XI during the mitotic cell cycle. **g,** Averaged contact signal of the 2µ plasmid in mutants depleted in Mcd1 (cohesin subunit), mutant depleted in Smc2 (condensin subunit) and in quiescence state.

H3 chemical cleavage data (Chereji et al., 2018) shows the regular distribution of nucleosomes along the genes, with nucleosome free regions at transcription start sites (TSS) and an even distribution along the open reading frames (**Fig. 1b**), showing that the chromatin of the plasmid is similar to that of its host. RNA-seq data highlight a relatively low transcriptional activity of the 4 genes (taking into account its size and the number of copies per cell, the level of transcription of 2µ plasmid is 10 times less than the average for its host), and the presence of non coding RNA as previously identified (Broach et al., 1979) (**Fig. 1b**). Finally, chromatin accessibility assessed by ATAC-seq confirms that the intergenic regions are the most open regions of 2µ plasmid (**Fig. 1b**). The cis-acting plasmid partitioning locus STB appears to be the most accessible region, which supports the notion that it acts as a gateway for known recruited host proteins (Chan et al., 2013). Note that no enrichment of the centromeric histone H3-like protein Cse4 was found on the plasmid sequence **(Supplementary Fig. 1**).

Overall, these signals confirm that the chromatin composition and organization of the 2µ plasmid is very similar to that of its host’s chromosomes (Nelson and Fangman, 1979).

### Discrete contacts between the 2µ plasmid and host chromosomes

To directly monitor the contacts made by the 2µ with the yeast genome, we plotted the relative contact frequencies between the plasmid and 16 yeast chromosomes from exponentially growing cells (Swygert et al., 2019) (**Fig. 1c, blue curve; Supplementary Fig. 2**). These curves can also be represented using a chromosomal heatmap diagram colored along its linear axis according to a scale that reflects contact frequencies (Methods; **Fig. 1c,d**, chromosomal shapes**; Methods**).

The 2µ plasmid contacts with the hosts chromosomes were not evenly distributed, as reflected by the curve peaks (dotted black boxes) and darker stripes along the chromosomal diagrams that represent hotspots of contacts (**Fig. 1c, black triangles**). A peak-calling algorithm yielded 73 such hotspots along the 16 host chromosomes, with most (66/73) located within chromosome arms, and 7 (more than randomly expected: 9.5% versus 3.9%) in subtelomeric regions (**Methods**). In contrast with past findings drawn from imaging approaches, centromeres were on average depleted in contact with the 2µ (**Fig. 1e**).

The contact pattern measured by Hi-C correlates very well with the Rep1 occupancy signal measured by ChIP-seq (**Supplementary Fig. 3**), supporting that the plasmid is indeed positioned in the close vicinity of discrete loci within the yeast genome. In contrast, a plasmid carrying a yeast centromeric sequence colocalizes near the SPB with the 16 yeast centromeres and displays contacts only with these regions (**Supplementary Fig. 4**a). A replicative plasmid devoid of centromere (pARS) displays relatively even contacts throughout the genome and does not show contact enrichment around the regions identified with the 2µ plasmid (**Fig. 1d, Supplementary Fig. 4**b). Altogether, these data show that the 2μ makes specific, discrete contacts with dozens of loci interspersed over the entire genome, excluding pericentromeric regions.

### REP1 and STB sequence are necessary for the specific attachment and stability

The dimer Rep1/Rep2 has been shown to associate with the plasmid STB sequence to promote partitioning during cell division and both STB and REP1/REP2 are required for the plasmid stability in host cell (McQuaid et al., 2019; Mereshchuk et al., 2022). We therefore tested whether the distribution of contacts along the genome was dependent on these partners by generating Hi-C data of cells with a 2µ plasmid mutants lacking either REP1 or the STB loci (Kikuchi, 1983; McQuaid et al., 2019) (**Methods**). These mutant plasmids are unstable (McQuaid et al., 2019) and carry a marker gene to be retained in the cell. This instability is reflected by a relatively low copy number per cell, which can be assessed from the proportion of reads from 2μ in each library (**Supplementary Fig. 5**).

In both mutants, there is no contact enrichment around the previously identified hotspots (**Supplementary Fig. 6 a,b**). Moreover, a plasmid mutant lacking only FLP1 shows contact enrichment around the identified hotspots (**Supplementary Fig. 6 c**) which indicates that this recombinase is not involved in establishing the specific contacts. These experiments show that the Rep1 protein and the STB sequence are essential for the establishment and/or maintenance of contacts between 2µ and specific regions of its host chromosomes.

### Plasmid/chromosomes contact regions are stable under a range of conditions

The diversity of published Hi-C and Micro-C experiments and the ubiquitous nature of the 2µ plasmid in yeast strains allowed sifting through existing data to detect variations in contact patterns between different genetic backgrounds, cell cycle stages, growth and metabolic states. Laboratory strains (W303 and S288C) and natural strains (BHB and Y9, (Peter et al., 2018)) displaying high or low copy number of plasmids respectively presented nearly identical contact hotspots profiles (**Supplementary Fig. 7**a,b) (**Methods**).

Furthermore, most hotspots were conserved throughout the mitotic cell cycle (**Fig. 1f**) (Costantino et al., 2020; Lazar-Stefanita et al., 2017) as well as during meiosis prophase (**Supplementary Fig. 8**) (Muller et al., 2017; Schalbetter et al., 2019). A small general increase in contact variability was observed at the later stages of meiosis prophase, which could reflect increased compaction and segregation of chromosomes into spores.

The hotspot pattern was also conserved upon the degradation of chromatin associated protein complexes including members of the structural maintenance of chromosome (SMC) family cohesin and condensin (**Fig. 1g**), known to organize host genome (Bastié et al., 2022; Dauban et al., 2020) and proposed to be involved in 2µ plasmid stability (Kumar et al., 2021); in cells lacking the silencing complex member Sir3 (**Supplementary Fig. 7**c) that is able to act as a bridging complex (Ruault et al., 2021); or in cells depleted for the DNA replication initiation factor Cdc45 that reach mitosis without replicating (Dauban et al., 2020) (**Supplementary Fig. 7**c). The pattern of interaction was also conserved in cells treated with nocodazole (**Supplementary Fig. 7**c). Previous reports pointed that nocodazole treatment impaired the recruitment of Cse4 (Hajra et al., 2006), Kip1 or microtubule associated proteins Bim1 and Bik1 (Prajapati et al., 2017) meaning that those host factors are not involved in the anchoring of the 2µ plasmid on host chromosomes.

The profile of contact of 2µ plasmid and host chromosomes appear very similar in other biological conditions like with double strand break (DSB) (Piazza et al., 2021), with hydroxyurea (HU) treatment (Jeppsson et al., 2022) and in different genetic mutants (**Supplementary Fig. 7**c).

In sharp contrast, most hotspots were strongly attenuated in quiescent cells (Guidi et al., 2015; Swygert et al., 2021) (**Fig. 1g**), when the cells dramatically alter their transcription program (McKnight et al., 2015) and genome organization (Guidi et al., 2015; Swygert et al., 2021). The later observation prompted us to explore more closely the links between transcription and the plasmid-chromosomal contacts.

### The 2µ preferentially tethers to inactive regions along the host’s chromosomes

To explore the links between transcription and plasmid contacts (**Fig. 2a)**, we first plotted the individual hotspots windows ordered by peak strength along with the corresponding transcription level and gene size annotation. This analysis reveals a contact enrichment with poorly transcribed regions often extending over several kbs (**Fig. 2b**), corresponding to relatively long genes (e.g. >4kb, to compare to a genomewide gene size median of 1kb) (**Fig. 2b**). They include 15 of the 19 genes longer than 7 kb that are non expressed in these growth conditions, with the remaining four being transcribed. A statistical analysis shows that 2µ plasmid binding depends on the level of transcription and the size of the gene (**Supplementary Fig. 9**). We piled-up the contacts made by the 2µ and 80 kb windows centered on the 73 peaks called on the contact profile, along with the pile-up transcription pattern of the 73 regions, revealing a strong depletion centered on the contact hotspot (**Fig. 2c; Methods**).

**Figure 2.**
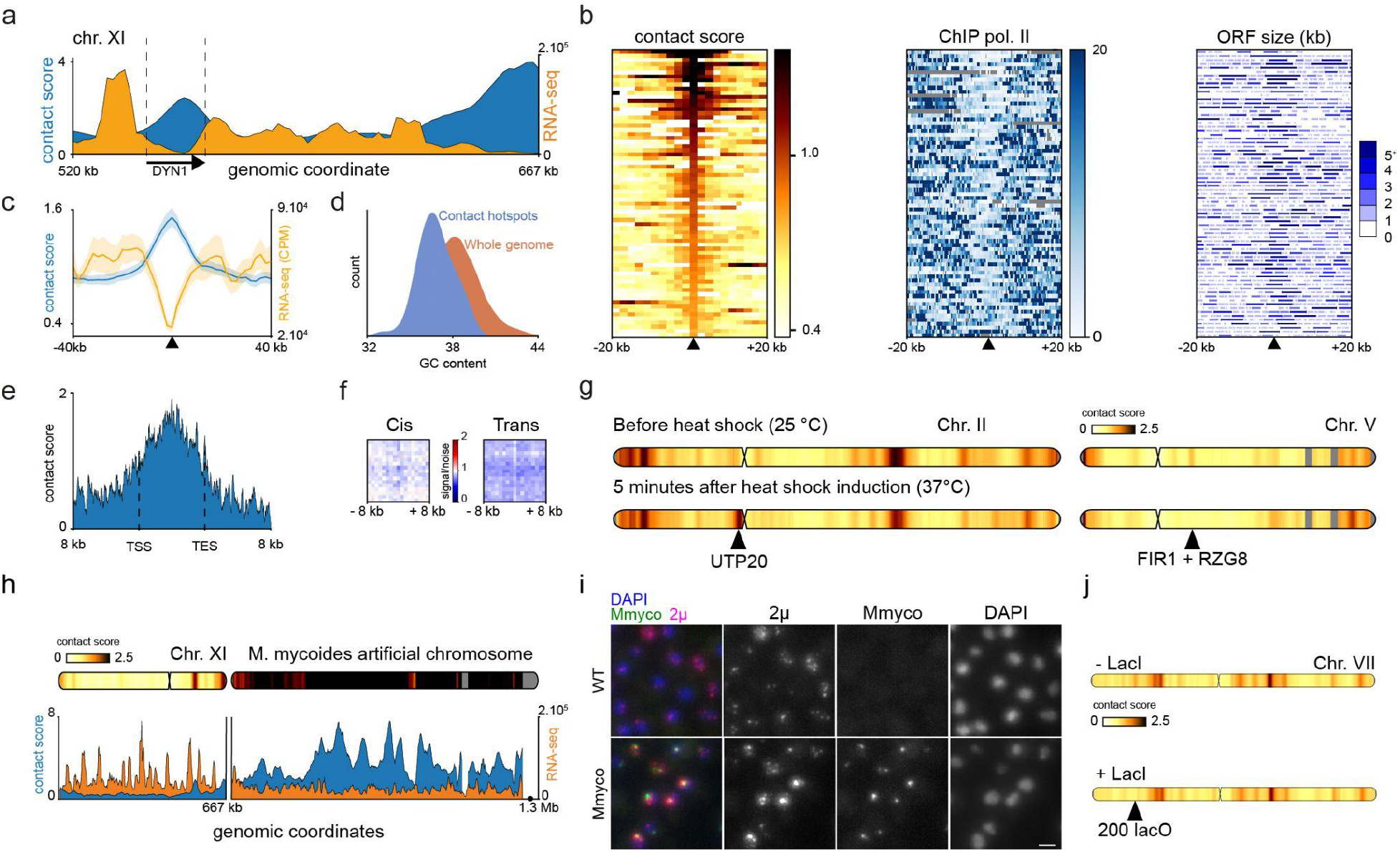
The contacted regions are depleted in transcription and more frequent at genes with long sizes. **a,** Transcription signal and contact profile of the 2µ plasmid in a region of the Chr. XI **b,** Heat maps of 2µ plasmid contact signals, transcription level (ChIP-seq of Pol II) and gene structure sorted in descending order according to contact scores over the region − 20 to + 20 kb around the peaks of contact of 2µ plasmid. **c,** Averaged contact signal of the 2µ plasmid and transcription at the positions of contact automatically detected in WT, log phase for 2µ. **d,** Distribution of GC content for the group of sequences contacted by the 2µ plasmid and for the whole genome of *S. cerevisiae*. **e,** Contact profile of 2µ plasmid along long genes (> 7 kb), binned at 200 bp. **f,** Mean profile heatmap between hotspots of contact with 2µ plasmid belonging to the same (left) or different chromosome (right). **g,** Contact profile of the 2µ plasmid before and 5 min after a heat shock for Chr. II and Chr. V. Examples of regions where contact intensity varies significantly are marked with a vertical arrow. **h,** Contact profile of the 2µ plasmid in strains containing an additional bacterial chromosome *M. mycoides* as well as the transcription profile. **i,** Representative fluorescent images (Z-stack projection) of FISH (Fluorescence In Situ Hybridization) experiments using a probe specific for the *M. mycoides* genome and a probe specific for the 2µ sequence. The probes were hybridized on a wild-type strain or a strain carrying the *M. mycoides* genome fused to chromosome XVI. The scale bar is 2µm. **j**, Contact profile of the 2µ plasmid at an array of 200 lacO binding sites without and with LacI protein. The region is marked with a vertical arrow head.

The average GC% of the hotspots sequences is slightly lower than the genome average (36.8% versus 38.2%) (**Fig. 2d)**, and no consensus was identified when processing hotspots sequences using MEME algorithm (Bailey et al., 2015) (**Methods**).

A magnification of contact distribution over the long inactive genes revealed a maximum enrichment in the middle of the sequence (e.g. DYN1 on **Fig. 2a**, **Fig. 2e**).

In addition, the regions contacted by 2µ are not enriched in *cis* or *trans* contacts with each other, suggesting they do not colocalize in the nuclear space (**Fig. 2f, Supplementary Fig. 10**a). Finally, the contact signal measured by Micro-C reveals an enrichment in short-range contacts at the hotspots (**Supplementary Fig. 10**b).

### Plasmid tethering is quickly reversible

We then explored the dynamics of plasmid chromosome anchoring. To do this, we induced heat shock stress, known to modify transcriptome, chromatin state and protein-genome interactions (Kim et al., 2010; Vinayachandran et al., 2018), by transferring exponentially growing cells at 25°C to 37°C medium (**Methods**).

Five minutes after heat shock, changes in the contact signal of 2µ plasmid were already observed. For instance, the contacts between the 2µ plasmid and the UTP20 gene (with a size of 7.4 kb) were strongly increased (**Fig. 2g**), while contact enrichment at the locus of FIR1 and RZG8 genes (with sizes of 2.6 kb and 3.2 kb) disappears (**Fig. 2g**). Precise kinetics with 4 time points show how contact points can appear (**Supplementary Fig. 11**a) or disappear (**Supplementary Fig. 11**b) in a matter of minutes. These results show that the plasmid is able to relocalize quickly to discrete regions.

### 2µ plasmid tether to exogenous artificial inactive chromatin

To further support the relationship between chromatin inactivity and 2μ contacts, we explored the plasmid behavior in presence of a Mb-long exogenous sequence. The plasmid positioning in strains carrying the linearized sequence of the *Mycoplasma mycoides* (Mmyco) chromosome as supernumerary, artificial chromosome, was investigated using Hi-C (Chapard et al., 2023) (Lartigue et al., 2009) (**Fig. 2h**). Mmyco presents highly divergent sequence composition, as reflected by its GC% (24% to compare with yeast 38%). It is chromatinized by well-formed nucleosomes, imposes little fitness cost to the yeast, and segregates properly. Mmyco AT-rich sequence is devoid of transcription and shows no enrichment in RNA Pol II (Chapard et al., 2023). Strikingly, contacts between the 2µ sequence and the entire length of Mmyco’s inactive 1.2Mb sequence were 6 fold higher than the average value on wild-type chromosomes. Our past work demonstrated that the Mmyco chromosome adopts a globular shape at the nuclear periphery (Chapard et al., 2023). In absence of Mmyco, the plasmid appears as several foci distributed in the nucleoplasm as previously reported (Heun et al., 2001; Velmurugan et al., 1998). In contrast, in the presence of the Mmyco, most of the 2µm signal concentrates and colocalizes with the bacterial chromosome., (**Fig. 2i**). This result demonstrates that the Hi-C data reflects titration of the 2µ plasmid from their hotspots by the long inactive sequence, and shows that inactivity is one of the primary conditions of plasmid relocalization to a sequence. In addition, this result also demonstrates that the contacts quantified using Hi-C between the 2µ plasmid and genomic DNA do indeed correspond to physical relocalization of the molecules.

We also analysed contacts between the 2µ and an artificial 9 kb array consisting of 200 lacO binding sites derived from *Escherichia coli* and introduced in chromosome VII (Guérin et al., 2019). The LacO array, which has a GC content of 41%, is not transcribed and is recognized by the DNA-binding repressor LacI put under the control of an inducible promoter (Guérin et al., 2019) (**Methods**). When LacI is not present, we observe a peak of contact but upon LacI binding, the array is not a contact hotspot anymore (**Fig. 2j**). These observations further support that a long (>9 kb) inactive region from a different organism but with a similar GC content than *S. cerevisiae* can be a contact hotspot for the 2µ plasmid. It has been shown that a high level of LacI binding results in nucleosome eviction (Loïodice et al., 2021). The observation that specific contact is lost when the LacI protein is attached to the region suggests that the resulting large nucleosome-free region could be responsible for the detachment of the 2µ plasmid.

### Characterisation of hot spots of contact of the 2μ plasmid

To further characterize the composition of the regions contacted by the 2μ plasmid, we computed the average enrichment of various genomic signals (ChIP-seq, MNAse-seq, ATAC-seq) over the 73 contact hotspots. For example, we computed the histone H3 ChIP-seq average signal and observed an enrichment at the hotspots (**Fig. 3a**). In agreement with the transcriptional inactivity of the hotspots, chromatin accessibility (ATAC-seq) (Lee et al., 2018) shows that these regions are less accessible compared to the rest of the host genome (**Fig. 3b**).

**Figure 3.**
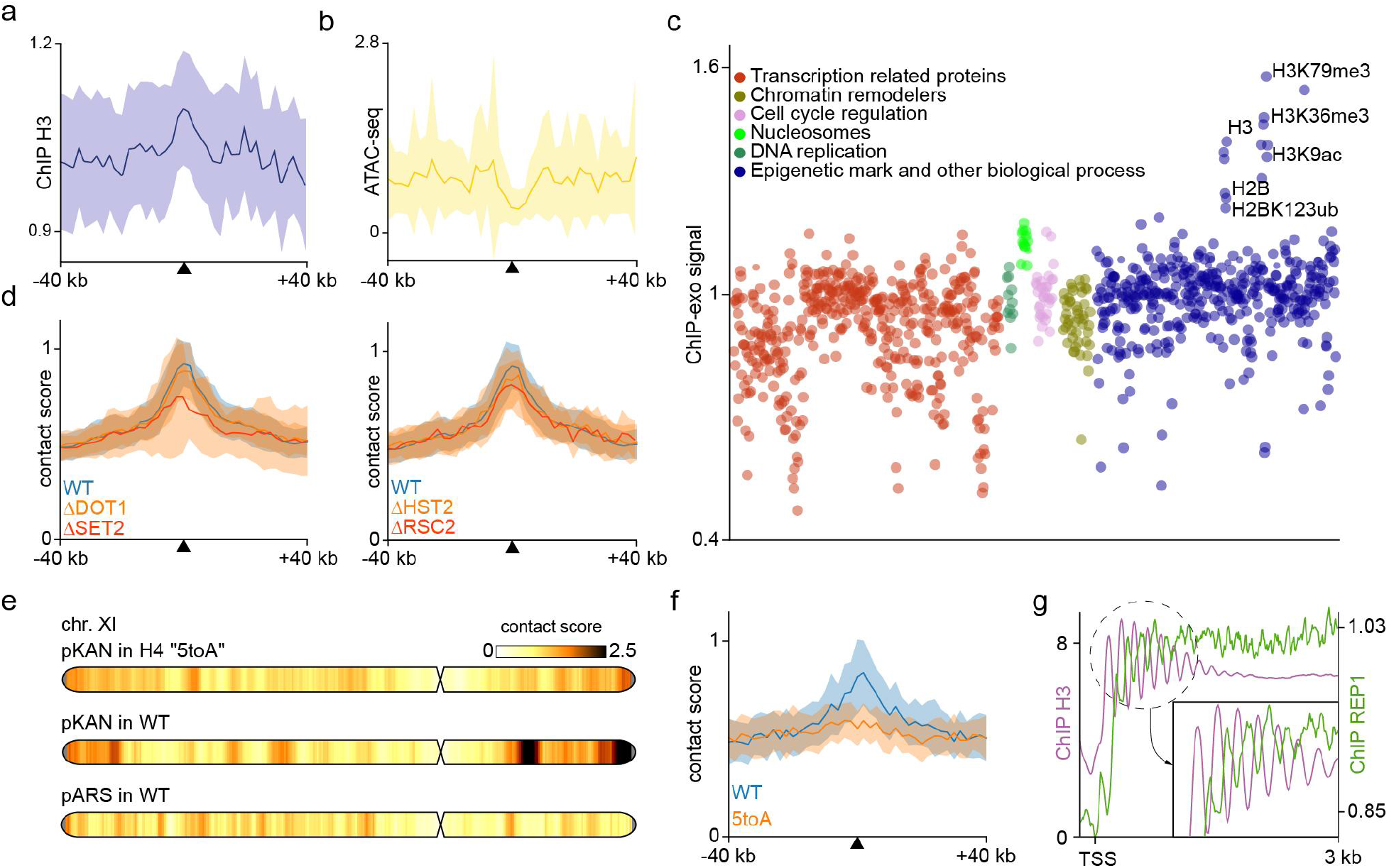
Specific positioning may be associated with nucleosome signal. **a,** Averaged signal at the hotspots of contact with 2µ plasmid for nucleosome occupancy (ChIP-seq of H3 histone) and **b**, chromatin accessibility (ATAC-seq). **c,** Average value of signal at the hotspots of contact for 1251 ChIP-exo libraries sorted by general categories (Rossi et al., 2021). **d,** Averaged contact signal of the 2µ plasmid in mutants of epigenetic marks ΔDot1, ΔSet2 and mutant in chromatin remodeler ΔRSC2 and in deacetylase mutant ΔHST2. **e,** Contact profile of the 2µ plasmid (pKAN version) with chromosomes of *S. cerevisiae* in H4 5toA mutant, WT and of the pARS plasmid in WT for chromosome XI. **f,** Averaged contact signal of the 2µ plasmid in H4 5toA mutant and control. **g,** Averaged signal around TSS for nucleosomes and REP1 occupancy signals.

To determine the chromatin composition of the chromosomal regions contacted by the 2µ plasmid, we also took advantage of a recently generated ChIP-exo dataset to screen for the deposition of ∼800 different proteins or histone marks along the *S. cerevisiae* genome (Rossi et al., 2021). For each genomic signal, the deposition profiles over the 73 contact hotspots were aggregated and tested for enrichment or depletion (**Methods**). In agreement with the low activity of the tested regions, most of the proteins associated with active transcription (e.g. general transcription factors or proteins of the SAGA complex) were depleted **(Fig. 3c**). On the other hand, histones H3, H2B, and histone marks like H3K79me3 and H3K36me3, were the only signals that were enriched over the contact hotspots (**Fig. 3c**).

We tested for the influence of both marks by characterizing the plasmid contacts in absence of either Set2 and Dot1 methylase. Absence of Dot1, responsible for H3K79 methylation, did not affect plasmid hotspots of contact (**Fig. 3d**). In absence of Set2, which methylates H3K36, long genes are known to be derepressed (Li et al., 2007). Although contact specificity for the 2µ plasmid is still detectable in this mutant (**Fig. 3d, Supplementary Fig. 12**b), long genes have their contact signal with the 2µ plasmid greatly reduced while the level of transcription is slightly increased (**Supplementary Fig. 12**c). Therefore, these experiments further confirm that transcription activity is linked to plasmid contact profile. In absence of silent regions, the plasmid appears to relocalize to the telomeric regions (**Supplementary Fig. 12**b). 2µ contact profile with the genome is also independent of remodeler complexes RSC2 or RSC1, and the centromere labeling protein HST2 (**Supplementary Figure 12d**). RSC2 was shown to be essential for the 2μm maintenance (Wong et al., 2002). We observed indeed a significant drop in the number of plasmids per cell in this mutant (**Supplementary Fig. 5**). However, its attachment to host chromosomes remain unchanged (**Supplementary Figure 12d**) suggesting that the mode of action of RSC2 is not directly linked to plasmid attachment. It was also shown for non-centromeric DNA circles that HST2 deacetylase was important for their condensation and propagation to daughter cells (Kruitwagen et al., 2018). However, we did not detect any changes in the contact profile of the HST2 deletion mutant (**Supplementary Figure 12d**). Finally, the 2µ contact profile with the genome is also independent of the main chromatin remodelers, as shown by the lack of variation that follows degradation using auxin-inducible degron (AID) of Spt6, Isw1, Swr1, Fun30, Ino80, Chd1, and Isw2 (Jo et al., 2021) (**Supplementary Figure 12d**).

### Plasmid anchoring depends on the H4 basic tail

A recent study suggested that chromatin folding at the nucleosomal level is altered when five basic amino acids of histone H4 tail (basic patches aa 15 to 20) are converted into alanine in a mutant called H4 5toA (Swygert et al., 2021). This change takes place without affecting transcription of the contact hotspots identified in WT condition (**Supplementary Fig. 13 a**). The 2µ plasmid contacts with chromosomes in the H4 5toA mutant were quantified using Hi-C (**Fig. 3e)**. No more contact enrichment on hotspots was detected (**Fig. 3f)**, suggesting that the 2µ plasmid contact hotspots depend on the presence or composition of the H4 tail basic patch. Importantly, in the H4 5toA mutant, transcription is not modified on the contact hotspots compared to WT condition (**Supplementary Fig. 13 a**). This result shows that the plasmid - chromosome contacts can be suppressed not only by transcription activation, but also only through an alteration of the nucleosome H4 tail basic patch. The same analysis was replicated in quiescent cells: in that condition, the remaining contact specificity between the 2µ plasmid and the 73 hotspots is also lost (**Supplementary Fig. 13 d**), suggesting the tail patch or more generally chromatin folding plays an important role in their maintenance.

A careful analysis showed that the Rep1 ChIP-seq signal is 90° phase-shifted with nucleosome position, (**Fig. 3g**) suggesting that Rep1 is not randomly contacting the host chromatin but is positioned in relation to the distribution of nucleosomes along the chromatin fiber.

Taken together, these results point to a model in which Rep1/Rep2 proteins localise to large regions that both are transcriptionally inactive and display a nucleosome signal carried by the H4 tail (**Fig. 4a**). Note that these two features could be two faces of the same coin since transcription disturbs nucleosome distributions, and could affect the histone signal specificity.

**Figure 4.**
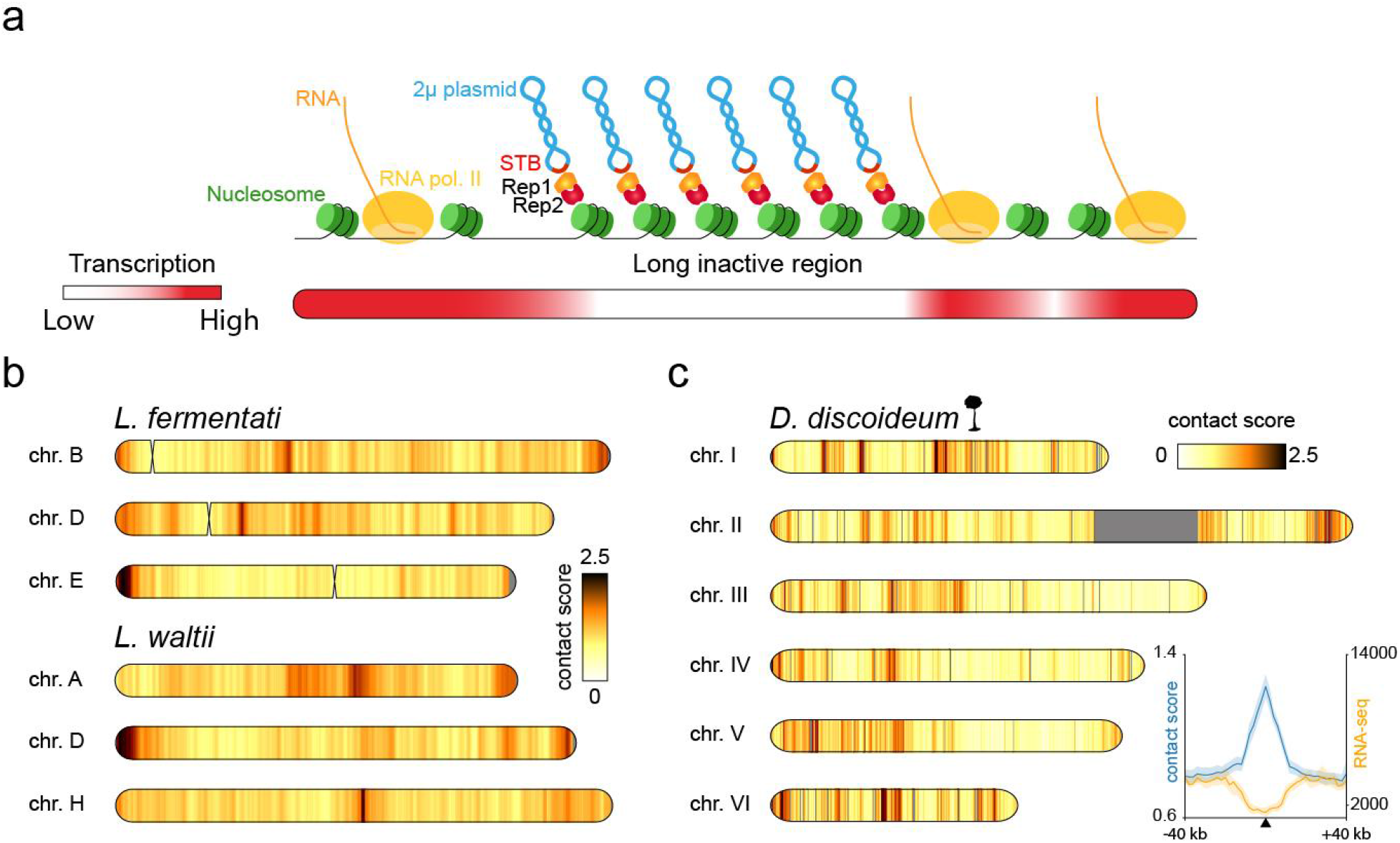
Contact specificity is also present in parasitic plasmids of other eukaryotes. **a,** Model proposed involving the plasmid proteins REP1/REP2 and nucleosome signal to make attachment between 2µ plasmid and specific loci on host chromosomes. **b** Contact profiles of natural plasmids with several chromosomes of the yeasts *Lachancea fermentati* and *Lachancea waltii.* **c,** Contact profile of the natural plasmid Ddp5 with chromosomes of the social amoeba *Dictyostelium discoideum.* Averaged contact and transcription signals around the loci detected as peaks of contact with Ddp5 plasmid.

### Other eukaryotic plasmids tethers to inactive chromatin

To assess whether this mechanism of binding to inactive sequences might concern other eukaryotic plasmids with ‬selfish” appearance, we analyzed contact profiles of *Lachancea fermentati* and *Lachancea waltii* which also host natural plasmids related to the 2µ plasmid. In these 2 organisms, we can also detect foci of contacts distributed across all the host chromosomes, far from the centromeres, and with also a bias towards long genes (**Fig. 4b, Supplementary Fig. 14**a).

We also quantified the contacts made by the Ddp5 plasmid with *Dictyostelium discoideum* chromosomes (Rieben et al., 1998). We performed Hi-C experiment on *D. discoideum* cells in vegetative state and measured the trans contacts between Ddp5 plasmid and its host chromosomes similarly to the 2μ plasmid experiment (**Fig. 4c, Supplementary Fig. 14**c,d). Around a hundred hotspots were detected on the contact profile (**Methods**). We computed the averaged transcription profile over windows containing these hotspots (Wang et al., 2021), the regions display reduced transcription compared to the rest of the genome (**Fig. 4c**), reminiscent of the 2µ plasmid hotspots.

## Discussion

In this work, we exploit overlooked genomic datasets to confirm that the 2μ plasmid chromatization is highly very similar to its host, potentially contributing to its long co-existence within its host’s nucleus (Lieberman, 2006). We also investigated using chromosome conformation capture data the physical contacts between the 2µ plasmid and its host’s chromosomes, revealing that the 2µ plasmid interacts to discrete positions along their arms. Most contact hotspots consist in long, poorly transcribed regions depleted in proteins of the transcription machinery or known DNA binding complexes, and might be associated with nucleosome signals as it depends on the tail of histone H4.

Previous reports have pointed out that the STB region on the 2µ plasmid recruits several yeast factors necessary for its segregation into the two daughter cells, including Cse4 (Hajra et al., 2006), the microtubule associated proteins Bim1 and Bik1, and the motor protein Kip1 (Prajapati et al., 2017). The disruption of microtubules using nocadazole led to the depletion of these proteins at STB region and generated plasmid missegregation (Prajapati et al., 2017), suggesting that those factors are important for the proper repartition between mother and daughter cells. However, HiC contact maps in presence of nocodazole treatment show that the plasmid remains bound to host chromosomes, indicating that these factors are not needed to maintain attachment. Moreover, condensin (Kumar et al., 2021) and cohesin (Mehta et al., 2002) were also proposed to be necessary for the plasmid propagation, but our data show that chromosome binding is independent of any both complexes. It cannot be ruled out that these proteins are useful to the 2µ plasmid but for other processes.

Observations obtained during heat shock experiments show that the kinetics of this system are of the order of a few minutes. These rapid kinetics are hardly compatible with processes for writing or erasing epigenetic marks. For example, it has been shown using optogenetic control of Set2 that the histone mark H3K36me3 has a writing and erasing time of around 30 minutes (Lerner et al., 2020). More generally, these experiments show that the attachment of the 2µ plasmid is dynamic and non specific to a DNA sequence, suggesting a high adaptability that can quickly adjust to the host metabolism.

Our experiments point to a model where the 2µ plasmid recognises through the REP1/REP2 proteins complex a structural signal involving several nucleosomes in the least active regions of its host chromosomes. The nature of this signal involves the basic tail of histone H4 but remains to be characterized . Since the 2µ plasmid binds to relatively long inactive regions, a possibility is that chromatin geometry plays a role in the attachments. For instance, the 2µ plasmid system may recognize specific chromatin fiber patterns, such as the alpha or beta nucleosome motifs recently characterized in Hi-CO (Ohno et al., 2019). The positioning along silent regions would allow it not to interfere with the biological processes of its host and thus to preserve a certain neutrality that may account for its low fitness costs and its long cohabitation with *S. cerevisiae*.

The present results presented here are also molecular experimental evidence in favor of the hitchhiking model (Sau et al., 2019) which proposes that the 2µ plasmid physically attaches to the chromosomes of its host in order to take advantage of all the machinery and chromosome movements during segregation. The probability of contact between two different molecules (inter chromosomal contacts) is very low from a thermodynamic point of view. The fact that we observe robust contact enrichment between the 2µ plasmid and the identified host positions is a strong indication that the plasmid must be physically attached to these positions.

Interestingly, we observed very similar behavior in other natural plasmids present in the nuclei of other eukaryotes: notably in the yeasts *L. waltii* and *L. fermentati*, but also in the amoeba *D. discoideum*, which is equidistant on the phylogenetic tree from the yeast *S. cerevisiae* and the human. The positioning characteristics of plasmid Ddp5 in *D. discoideum* are very similar to those we have demonstrated in *S. cerevisiae* (long non-transcribed regions). This suggests that the mechanism used may be the same between these different organisms. Whereas the CRISPR-cas9 system is based on a recognition mechanism that relies on a precise DNA sequence, the host-parasite system studied here seems to reveal a specificity mechanism based on a structural signal involving several nucleosomes. We envision that other biological processes depend on information encoded in nucleosomal availability and/or chromatin folding notably chromosome attachment of certain DNA viral episomes like Epstein Barr virus (EBV) (Kim et al., 2020) or Kaposi’s sarcoma-associated herpesvirus (KSHV) or other extrachromosomal circular DNA (eccDNA) (Møller et al., 2015).

## Acknowledgements

The authors thank M. Dobson for sharing 2µ plasmid mutant strains and Rep1 antibody. We thank G. Liti and J. Schacherer for sharing strains and suggestions. We thank S G Swygert and T Tsukiyama for sharing strains, Y. Barral, L. Baudry, G. Millot, P. Moreau, A. Piazza for useful discussions; C. Chapard, C. Matthey-Doret, J. Sérizay for technical advice. This work used the computational and storage services (maestro cluster) provided by the IT department at Institut Pasteur, Paris. F.G is supported by an ENS Paris Saclay fellowship. This work has received support under the program Investissements d’Avenir launched by the French Government and implemented by ANR with the references ANR–10–LABX–54 MEMOLIFE and ANR–10–IDEX–0001–02 PSL Université Paris, Q-life ANR-17-CONV-6150005 (SA). The authors greatly acknowledge the PICT@Pasteur imaging facility of the Institut Curie, member of the France Bioimaging National Infrastructure (ANR-10-INBS-04). This research was supported by funding from the European Research Council under the Horizon 2020 Program (ERC grant agreement 771813) to R.K, from Agence Nationale pour la Recherche ANR-22-CE12-0013-01 to R.K. and A.T., and from Agence Nationale pour la Recherche JCJC 2019 (Apollo) to A.C.

## Author contributions

All authors read and approved the final manuscript.

## Competing financial interests

The authors declare no competing financial interests.

## Material and Methods

### Strains and medium culture conditions

The genotype and background of strains used in this study are listed in the strain table (**Supplementary Table 3**). The jhd2::KANMX4, set2::KANMX4, dot1::KANMX4, rsc1::KANMX4, rsc2::KANMX4, hst2::KANMX4 strains were made using PCR amplified regions of strain from EUROSCARF collection. The diploid strain containing Mycoplasma chromosomes were made by crossing a strain containing a CRISPR linearized version of Mycoplasma chromosomes with BY4742.

Culture media Liquid YPD media (1% Yeast extract, 2% peptone, 2% Glucose), containing 200 µg.mL-1 of Geneticin (Thermofisher CAT11811031) or not, and SD-HIS (0.17% Yeast Nitrogen Base, 0.5% Am-monium Sulfate,0.2% synthetic dropout lacking histidine and 2% Glucose) were prepared according to standard protocols.

*Dictyostelium discoideum* cells were cultured in 20 ml autoclaved SM medium (per L: 10g glucose, 10g proteose peptone, 1g yeast extract, 1g MgSO_4_*7H_2_O, 1.9g KH_2_PO_4_, 0.6g K_2_HPO_4_) with dead *Klebsellia pneumoniae* at 20°C and 130 rpm. After 4 days of growth, cells were centrifuged at 300 rpm during 10 min before performing Hi-C procedure.

### Heat-shock experiment

Fresh YPD media was inoculated with overnight culture of C+ BY4741 cells (YPD, 25°C) and grown at 25°C. When the culture reached 10^7^ cells.mL^-1^, heat shock was applied by adding warm (65°C) fresh YPD media in order to shift media temperature from 25°C to 37°C. Cells were grown at 37°C and cells were extracted at different timepoints for Hi-C and ChIP-seq.

### ChIP-seq procedure

ChIP was performed as described previously (Hu et al., 2015) without calibration strain. 15 OD_600_ unit of *S. cerevisiae* (approximately 1,5.10^8^ cells) were harvested from an exponentially growing culture. Cell lysis was performed using Precellys in 2mL VK05 tubes and the sonication was performed on a Covaris S220 system as described previously (Piazza et al., 2021). To pull down Rep1 protein we used a polyclonal antibody, production has been previously described (Sengupta et al., 2001). Pull down chromatin was purified and prepared for paired end sequencing as described previously (Bastié et al., 2022).

### FISH

FISH (Fluorescence In Situ Hybridization) experiments were performed as in (Gotta et al., 1996) with some modifications. The *M. mycoides* probe was obtained by direct labeling of the bacterial DNA (1.5 µg) using the Nick Translation kit from Jena Bioscience (Atto488 NT Labeling Kit). For the 2µ plasmid labeling, a 5-kb PCR fragment was amplified from the 2µ plasmid DNA using primer pair FG92 (TTTCTCGGGCAATCTTCCTA) / FG24 (GTATGCGCAATCCACATCGG). This PCR product (1.5 µg) was then labeled using the Nick Translation kit from Jena Bioscience (AF555 NT Labeling Kit). For the *M. mycoides* probe and the 2µ plasmid probe, the labeling reaction was performed at 15°C for 90 min and 30 min, respectively. The labeled DNA was purified using the Qiaquick PCR purification kit from Qiagen, eluted in 30 µl of water. The purified probe was then diluted in the probe mix buffer (50% formamide, 10% dextran sulfate, 2× SSC final). 20 OD of cells (1 OD corresponding to 10^7^ cells) were grown to mid–logarithmic phase (1–2 × 10^7^ cells/ml) and harvested at 1,200 *g* for 5 min at RT. Cells were fixed in 20 ml of 4% paraformaldehyde for 20 min at RT, washed twice in water, and resuspended in 2 ml of 0.1 M EDTA-KOH pH 8.0, 10 mM DTT for 10 min at 30°C with gentle agitation. Cells were then collected at 800 *g*, and the pellet was carefully resuspended in 2 ml YPD - 1.2 M sorbitol. Next, cells were spheroplasted at 30°C for 10 minutes with Zymolyase (60 µg/ml Zymolyase-100T to 1 ml YPD-sorbitol cell suspension). Spheroplasting was stopped by the addition of 40 ml YPD - 1.2 M sorbitol. Cells were washed twice in YPD - 1.2 M sorbitol, and the pellet was resuspended in 1 ml YPD. Cells were put on diagnostic microscope slides and superficially air dried for 2 min. The slides were plunged in methanol at -20°C for 6 min, transferred to acetone at -20°C for 30 s, and air dried for 3 min. After an overnight incubation at RT in 4× SSC, 0.1% Tween, and 20 μg/ml RNase, the slides were washed in H_2_0 and dehydrated in ethanol 70%, 80%, 90%, and 100% consecutively at -20°C for 1 min in each bath. Slides were air dried, and a solution of 2× SSC and 70% formamide was added for 5 min at 72°C. After a second step of dehydration, the denatured probes were added to the slides for 10 min at 72°C followed by a 37°C incubation for 24h in a humid chamber. The slides were then washed twice in 0.05× SSC at 40°C for 5 min and incubated twice in BT buffer (0.15 M NaHCO_3_, 0.1% Tween, 0.05% BSA) for 30 min at 37°C. For the DAPI staining, the slides were incubated in a DAPI solution (1µg/ml in 1× PBS) for 5 minutes and then washed twice in 1× PBS without DAPI.

### Microscopy and image analysis

For all fluorescent images, the axial (z) step is 200 nm and images shown are a maximum intensity projection of z-stack images. Images were acquired on a wide-field microscopy system based on an inverted microscope (TE2000; Nikon) equipped with a 100/1.4 NA immersion objective, a C-mos camera and a Spectra X light engine lamp (Lumencor, Inc) for illumination. The microscope is driven by the MetaMorph software (Molecular Devices). Images were not processed after acquisition. Images shown are maximum intensity projection of Z-stack acquisition.

### Hi-C procedure and sequencing

Cell fixation was performed with 3% Formaldehyde (Sigma-Aldrich cat no F8775) and performed as described previously (Dauban et al., 2020). Quenching of formaldehyde was done by adding 300 mM of glycine at room temperature for 20 mins. All of the Hi-C were done using Arima Hi-C kit (Arima Genomics; restriction enzymes: DpnII, HinfI). Sequencing preparation was done using a Colibri ES DNA Library Prep kit for Illumina Systems (A38606024) and then sequenced on Illumina NextSeq500.

The 2 Hi-C libraries concerning *Lachancea waltii* and *Lachancea fermentati* were generated using a different Hi-C protocol described previously (Lazar-Stefanita et al., 2017).

## Data analysis

### Contact data processing (Hi-C, Micro-C)

Hi-C and MicroC processing was performed using hicstuff package (Matthey-Doret et al., 2022). Briefly, the paired-end reads were aligned to the S288C reference genome (GCA-000146045.2, R64-1-1) and the 2µ plasmid sequence (GenBank accession number: CM007980) as well as with *M. mycoides* (GCA-006265075). For the two experiments containing the lacO site array, reads were aligned to genomes based on strain W303 containing the lacO site sequences (for details on the constructions, see (Guérin et al., 2019)). A threshold of 1 for mapping quality was used for these 2 experiments. Genomes for *Lachancea waltii* was CBS6430 and X56553.1 sequence was used for pKW1 plasmid. Genome for *Lachancea fermentati* was CBS 6772 and M18275.1 sequence was used for the plasmid pSM1. For *Dictyostelium discoideum*, AX4 reference genome sequence GCA_000004695.1 dicty_2.7 was used and NC_001889.1 sequence was used for Ddp5 plasmid.

We used bowtie2 in its very sensitive local mode (Langmead and Salzberg, 2012). Unique mapped paired reads are then assigned to restriction fragments and non-informative contacts are filtered (Cournac et al., 2012; Matthey-Doret et al., 2022). PCR/optical duplicates are discarded (i.e paired reads mapping at exactly the same genomic positions). Contact signals were binned at 2 kb resolution except where noted (e.g. for the 2µ plasmid contact map or the averaged contact signal at long genes, a resolution of 200 bp was used).

### Computation of contact signal of 2µ plasmid

To compute the contact signal of the 2µ plasmid with each bin of the host genome, we used the normalized following score:

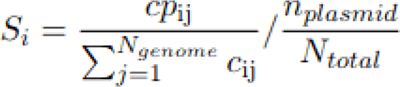

S_i_ represents the contact score between 2µ plasmid and a bin i in the host genome. It corresponds to the proportion of contacts made with the 2µ plasmid for a bin i of the host genome normalized by the percentage of presence of 2µ plasmid in the library. cp_ij_ is the number of contacts detected between 2µ plasmid and host bin i. 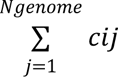 is the total number of contacts detected for the host bin i. *N_genome_* is the total number of genomic bins (in general 2 kb bins). *n_plasmid_* is the number of reads involving the plasmid. *N_total_* is the total number of reads of the library. For idiogram visualization, the contact signal S_i_ is represented using R ideogram package (Hao et al., 2020).

### ChIP-Seq, Mnase-seq, ATAC-seq and RNA-seq processing

ChIP-Seq, Mnase-seq, ATAC-seq and RNA-seq processing was performed using TinyMapper (https://github.com/js2264/tinyMapper). Paired end reads were aligned using the very sensitive and local mode of bowtie2 (Langmead and Salzberg, 2012), against S288C reference genome and the 2µ plasmid (CM007980) sequences . Only concordant pairs were retained, and reads with a mapping quality larger than 0 were kept. PCR duplicates were removed using samtools. When available, the input was similarly processed. Coverage for each genomic position was computed using deeptools bamcoverage function then normalized by Count Per Million (CPM) method. ChIP signal was then computed by dividing immuno-precipited normalized coverage by input coverage. Signals were then visualized with home made python using pyBigWig package. RNA-seq signal was log represented.

### Automatic detection of peaks of contact with 2µ plasmid

To detect contact peaks with the 2µ plasmid, we used the *find_peaks* function of the scipy package (with the following parameters: height = 0.8, distance=2). Before detection, the contact signal was interpolated using the *interp1d* function of the scipy package. In most of the average 2µ plasmid contact profiles shown, the set of contact peaks used is the one detected under normal log-phase culture conditions from Micro-C data (Swygert et al., 2019) corresponding to the 73 genomic positions given in **Extended Table 1**.

### Averaged contact profile of 2µ plasmid around loci

To plot the average contact profile around loci of interest, we extracted the contact signals at windows +/- 40 kb centered at positions of interest using a 2 kb binned signal. In cases where the limits of the window exceed a chromosome, Nan values were used. The standard deviation is represented around the mean value.

### Genomic features of regions contacted by 2µ plasmid

To plot the aggregated profile of genomic signals, we convert contact data (cool file at 200 bp resolution) into a bw file using home made python code. We then used the functions *computeMatrix* and *plotProfile* from deepTools suite (Ramírez et al., 2016) to compute and plot the heatmaps. Same approach was used to plot the averaged contact signal inside long genes. Gene boxes at regions contacted by 2µ plasmid were generated using home made python code and using the coordinates of genes of SGD database (http://sgd-archive.yeastgenome.org/sequence/S288C_reference/orf_dna/).

To plot the pileup plots around the pairs of genomic positions of peaks of contact with 2µ plasmid, we used the *quantify* mode of Chromosight (Matthey-Doret et al., 2020) with the following parameters --perc-undetected=100 --perc-zero=100. All possible pairs of peaks of contact in intra or inter configurations were generated using home made python code and the function *combinations* from itertools package.

The computational screen uses a dataset of Chip-exo libraries for about 800 different proteins and genomic signals (Rossi et al., 2021). For each aligned library (bowtie2 alignment with very sensitive mode and mapping quality >0), Chip-exo signal was computed with home made python code with a binning of 2 kb. An enrichment score was computed by taking the average of the ChIP-exo signal +/- 2kb signal around the bins of the positions of contact peaks detected in log phase (**Extended Table 1**). The different libraries were sorted according to the different categories identified in the UMAP analysis of (Rossi et al., 2021).

## Data availability

Some of the data associated with this study are publicly available and their reference numbers are listed in Supplementary Tables 1 and 2.

## Code availability

All scripts required to reproduce figures and analyses are available at https://github.com/acournac/2micron-project.

## Supplemental information

### Anchoring of parasitic plasmids to inactive regions of eukaryotic chromosomes through nucleosome signal

**Supplementary Figure 1.**
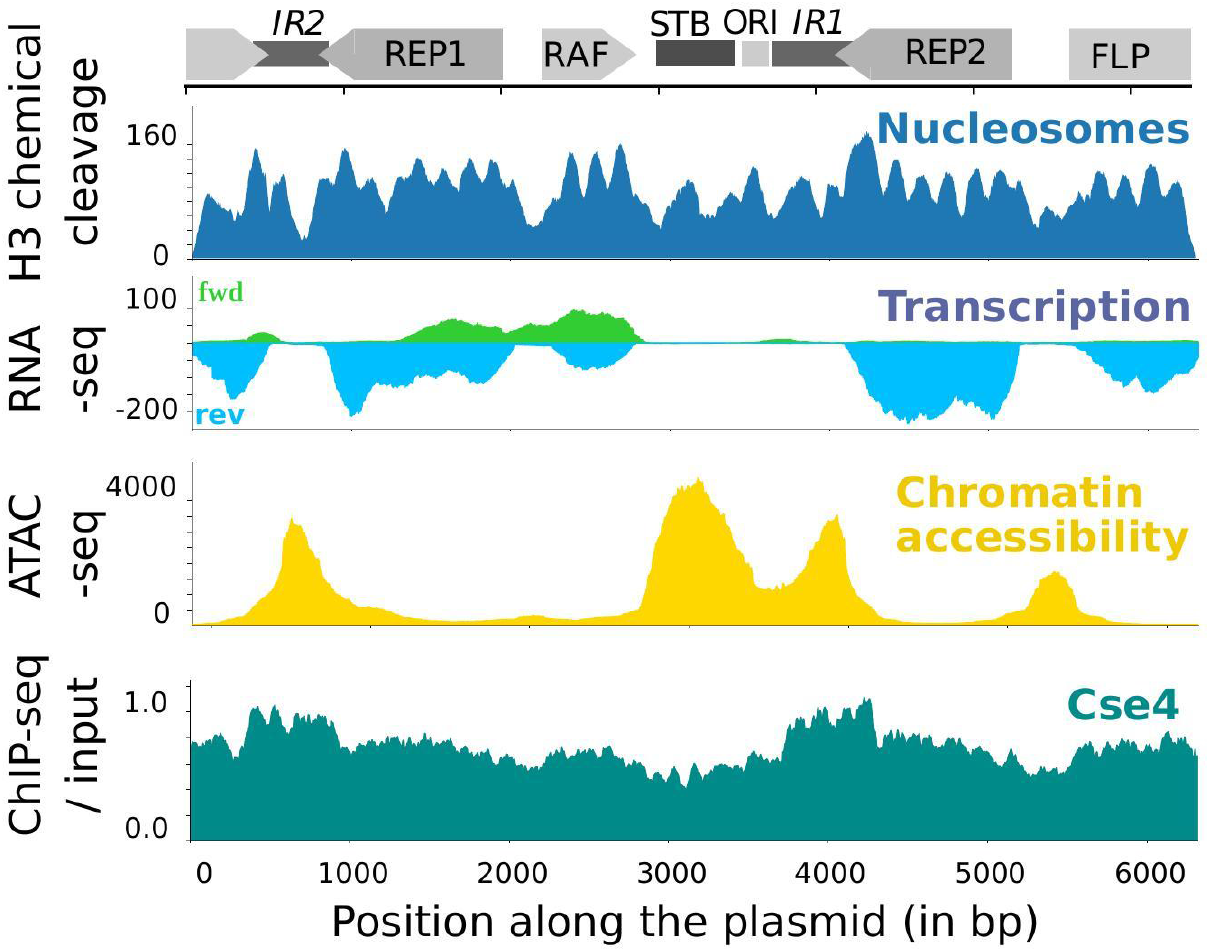
Genomic signals along the 2*µ* plasmid of *Saccharomyces cerevisiae*. Nucleosomes signal along the 2*µ* plasmid from H3 chemical cleavage data [1]. Transcription signal along the 2*µ* plasmid from RNA-seq data of [2]. Chromatin accessibility signal along the 2*µ* plasmid from ATAC-seq data of [3]. Protein occupancy of Cse4 along the 2*µ* plasmid, from ChIP-seq data of [4].

**Supplementary Figure 2.**
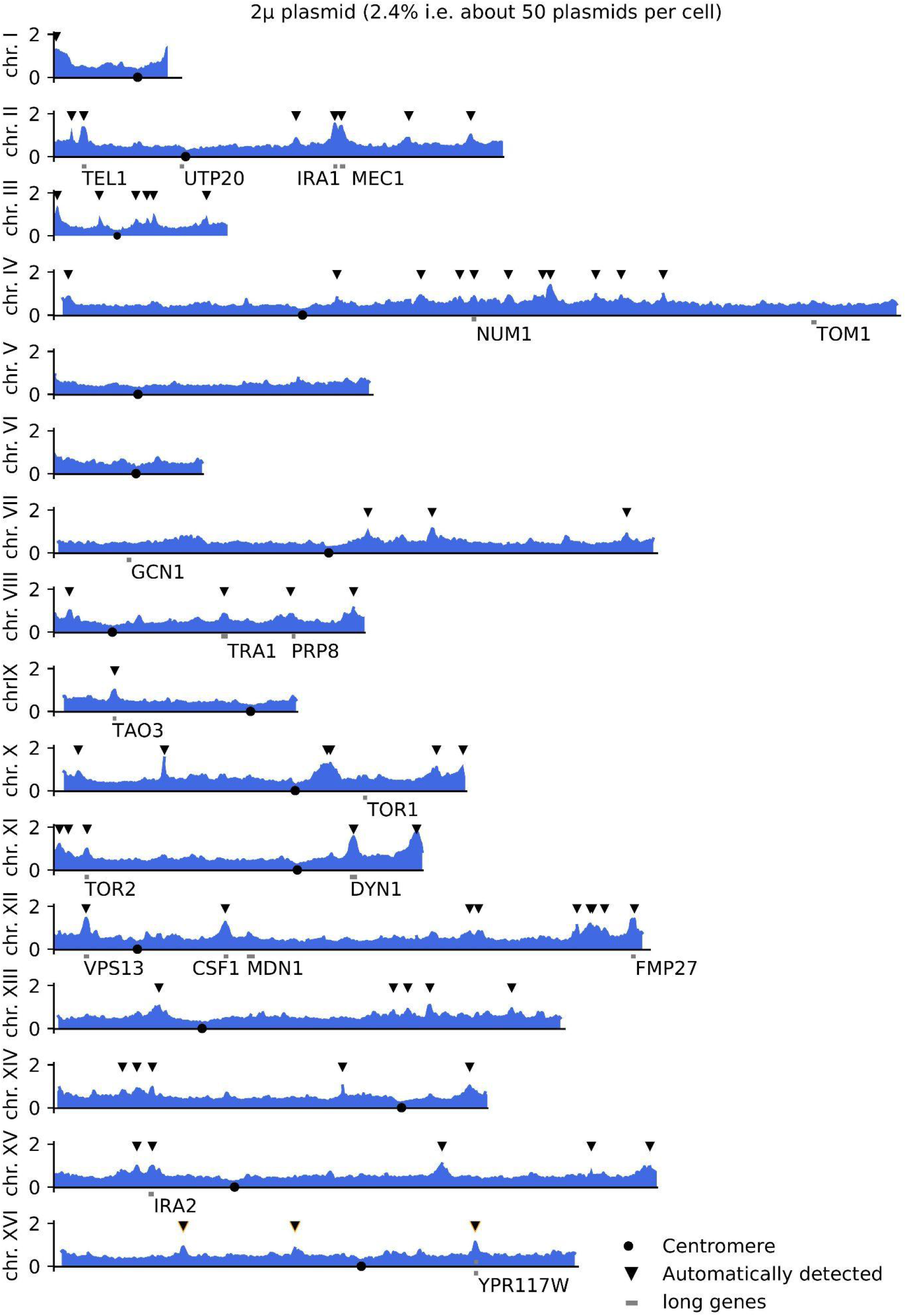
Contact signal of the 2*µ* plasmid along the 16 chromosomes of S. cerevisiae. **a**, The contact signal is binned at 2 kb, genes with size *>* 7 kb are annotated with grey rectangles and their names, (MicroC data from [5]). Automatically detected peaks of contact were annotated with black triangles.

**Supplementary Figure 3.**
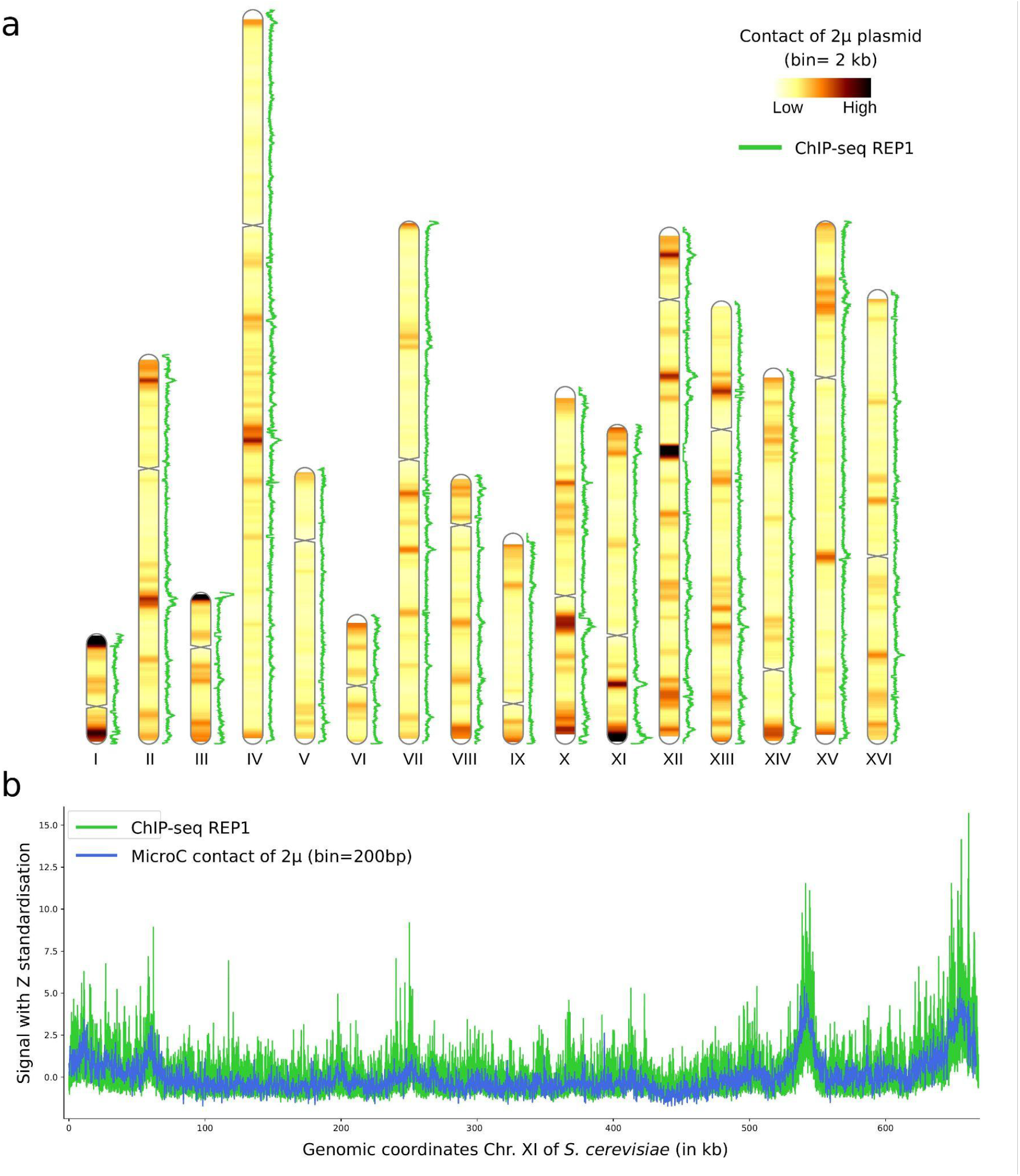
ChIP-seq of REP1 protein from the 2*µ* plasmid. **a**, ChIP-seq of Rep1 protein along with the contact profile of 2*µ* plasmid with the chromosomes of *S. cerevisiae* (chromosomal heatmap diagram). **b**, ChIP-seq of Rep1 protein and contact profile of 2*µ* plasmid binned at 200 bp, (MicroC data from [5]) for the chromosome XI of *S. cerevisiae*.

**Supplementary Figure 4.**
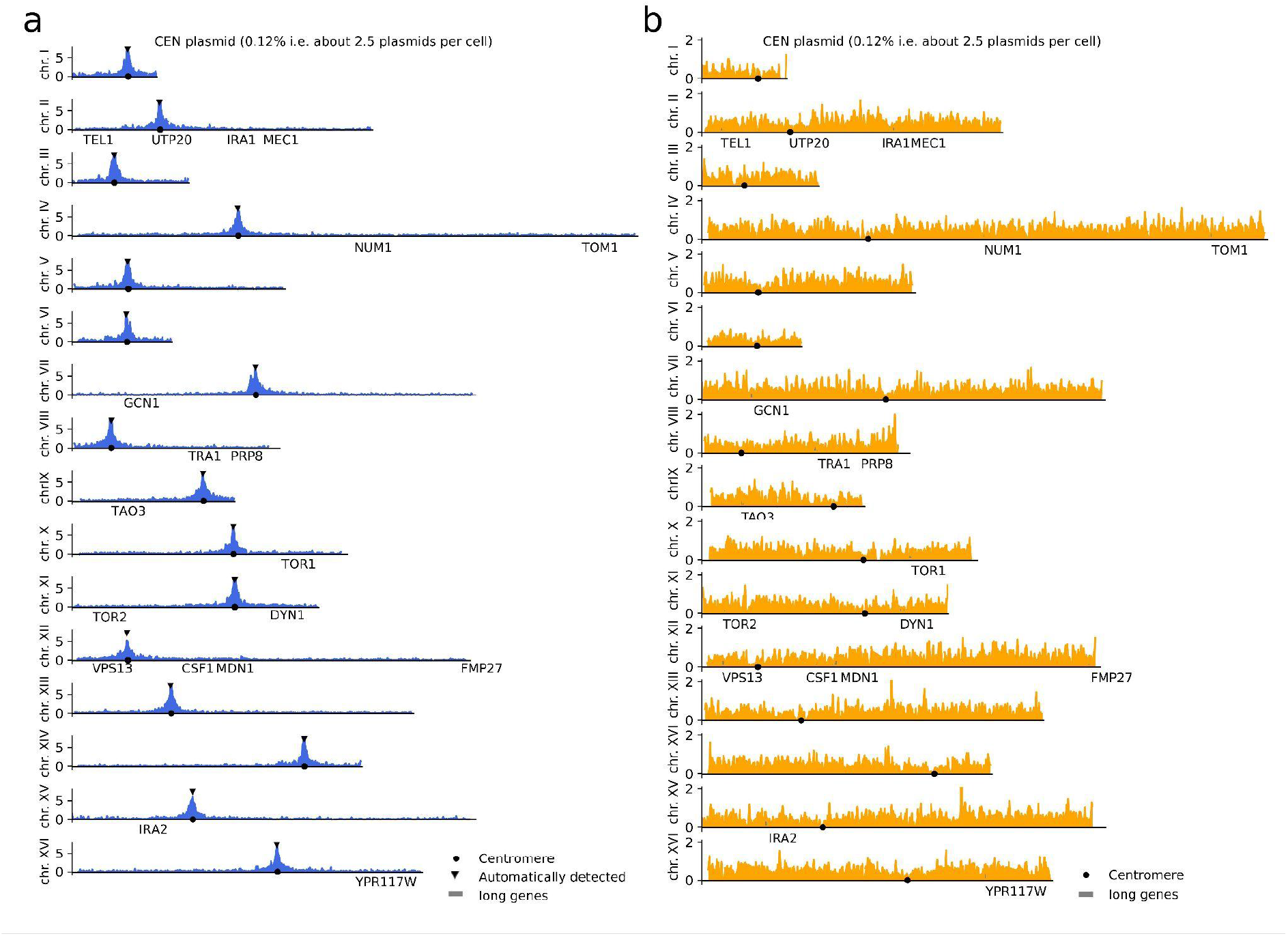
Contact signal of the control plasmids along the 16 chromosomes of *S. cerevisiae*. **a,** Contact signal of the yeast centromeric Plasmid (YCp) pRS416 along the 16 chromosomes of *S. cerevisiae*. The Hi-C contact signal is binned at 2 kb, names of genes with size *>*7 kb are annotated. Automatically detected peaks of contact were annotated with black triangles. **b,** Contact signal of a replicative plasmid devoid of centromere (pARS) and 2*µ* system along the 16 chromosomes of *S. cerevisiae*. The contact signal is binned at 2 kb, names of genes with size *>*7 kb are annotated.

**Supplementary Figure 5.**
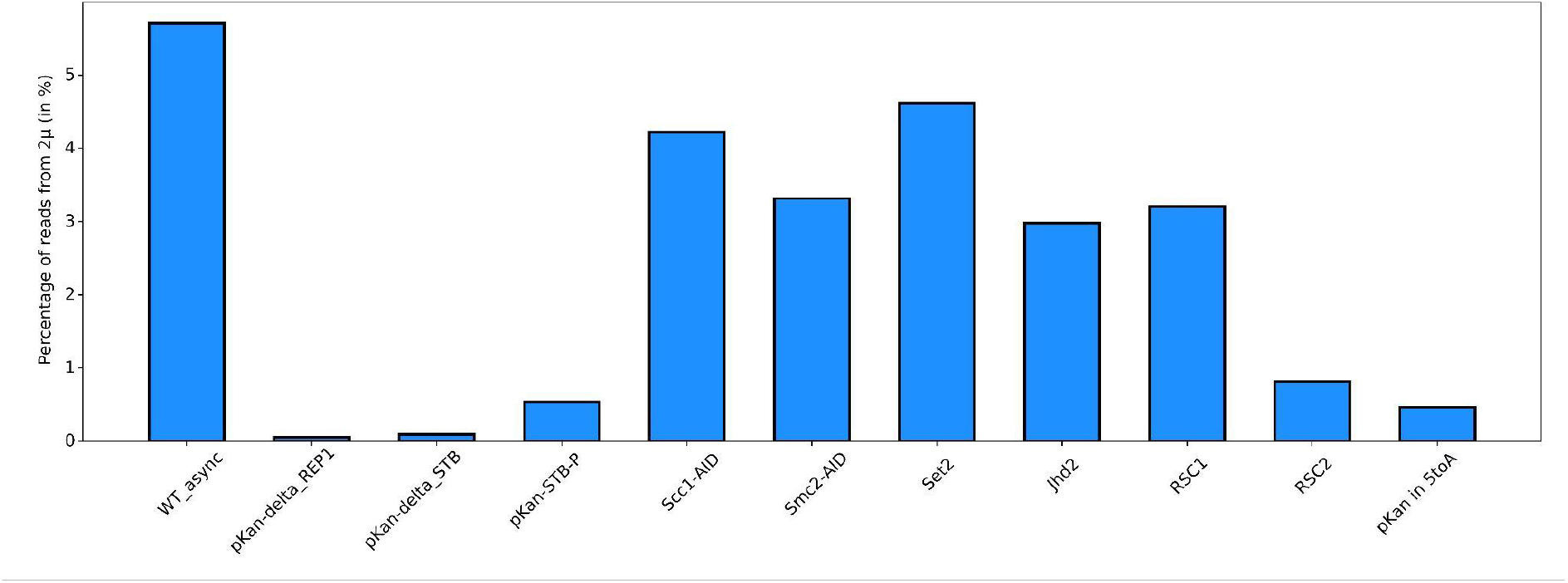
Percentage of reads coming from 2μ plasmid sequence in WT and various mutants.

**Supplementary Figure 6.**
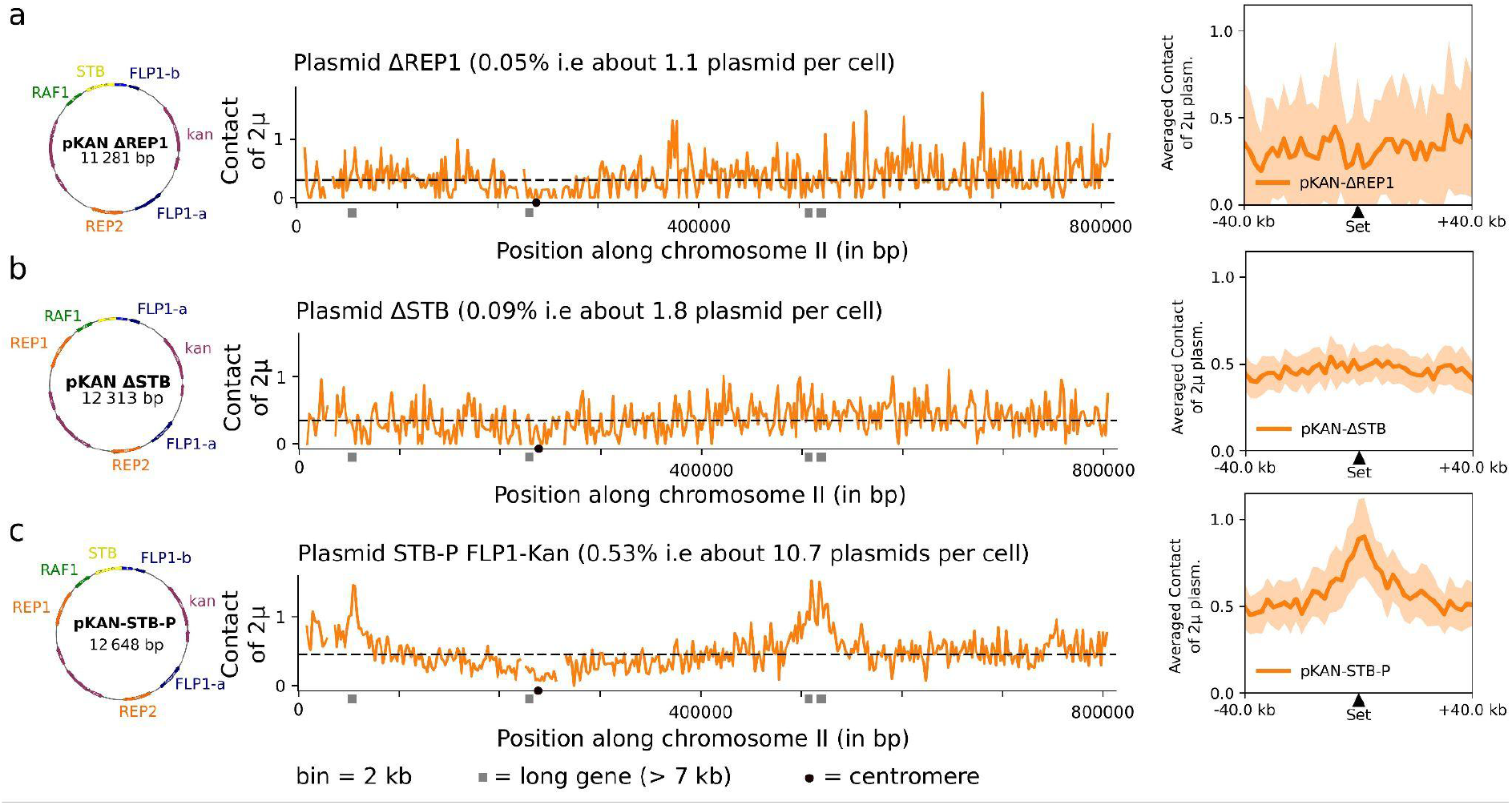
Contact signal of 2*µ* plasmid mutants. **a**, Contact signal of the ΔREP1 mutant 2*µ* plasmid along the chromosome II of *S. cerevisiae* and the averaged contact signal on the hot spots of contact detected in WT, log phase condition. **b**, Same for the ΔSTB mutant 2*µ* plasmid. **c**, Same for the STB-P 2*µ* mutant plasmid.

**Supplementary Figure 7.**
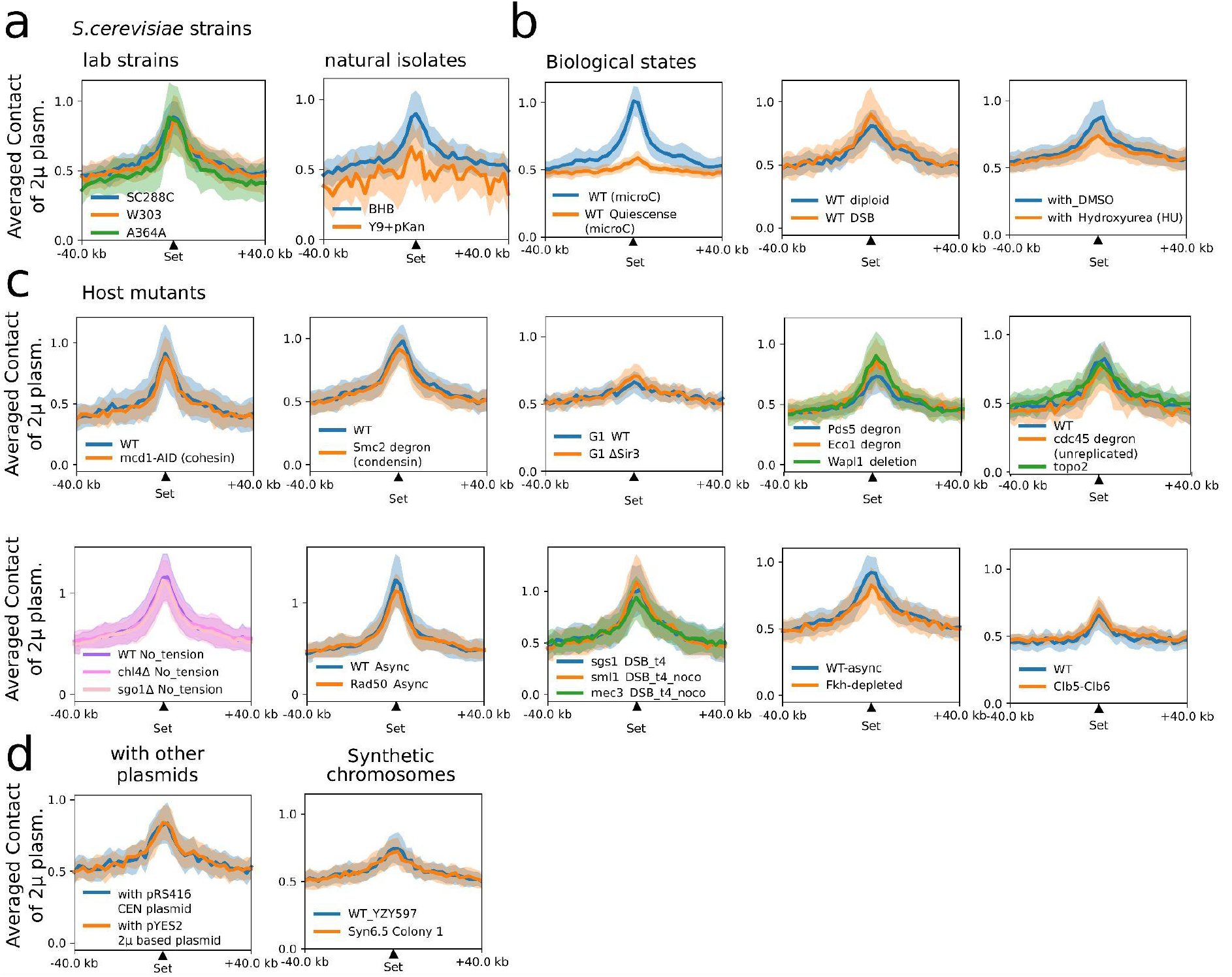
The specific positioning of the 2*µ* plasmid is conserved under a wide variety of biological conditions and mutants. **a**, Averaged 2*µ* plasmid contact signal over the hotspots of contact identified in WT, log phase in different lab strains of *S. cevevisiae*: SC288C, W303 [6], A364A [7] and strains from natural isolates [8]. **b**, Averaged 2*µ* plasmid contact signal over the hotspots of contact identified in WT, log phase in different biological states: in quiescence [9], in diploid stage, with double strand break of DNA [10], with DMSO or HU treatment [11]. **c**, Averaged 2*µ* plasmid contact signal over the hotspots of contact identified in WT, log phase in different mutants of *S. cevevisiae*: Mcd1 depleted (sub-unit of cohesin) [7], Smc2 depleted (sub-unit of condensin) [12], ΔSir3 [9], Pds5, EcoI, Wapl [6], in cdc-45 degron mutant (stopped replication) [6], ΔTOP2 [13], in condition with no tension of microtubules (nocodazole treatment), ΔChl4, ΔSgo1 [14], ΔRad50 [15], ΔSgs1, ΔSml1, ΔMec3 [10], in Fkh-depleted mutant [16], in Clb5-Clb6 mutant [17]. **d**, Averaged 2*µ* plasmid contact signal over the hotspots of contact identified in WT, log phase in presence of other plasmids (centromeric and 2µ based) and with synthetic chromosomes [18].

**Supplementary Figure 8.**
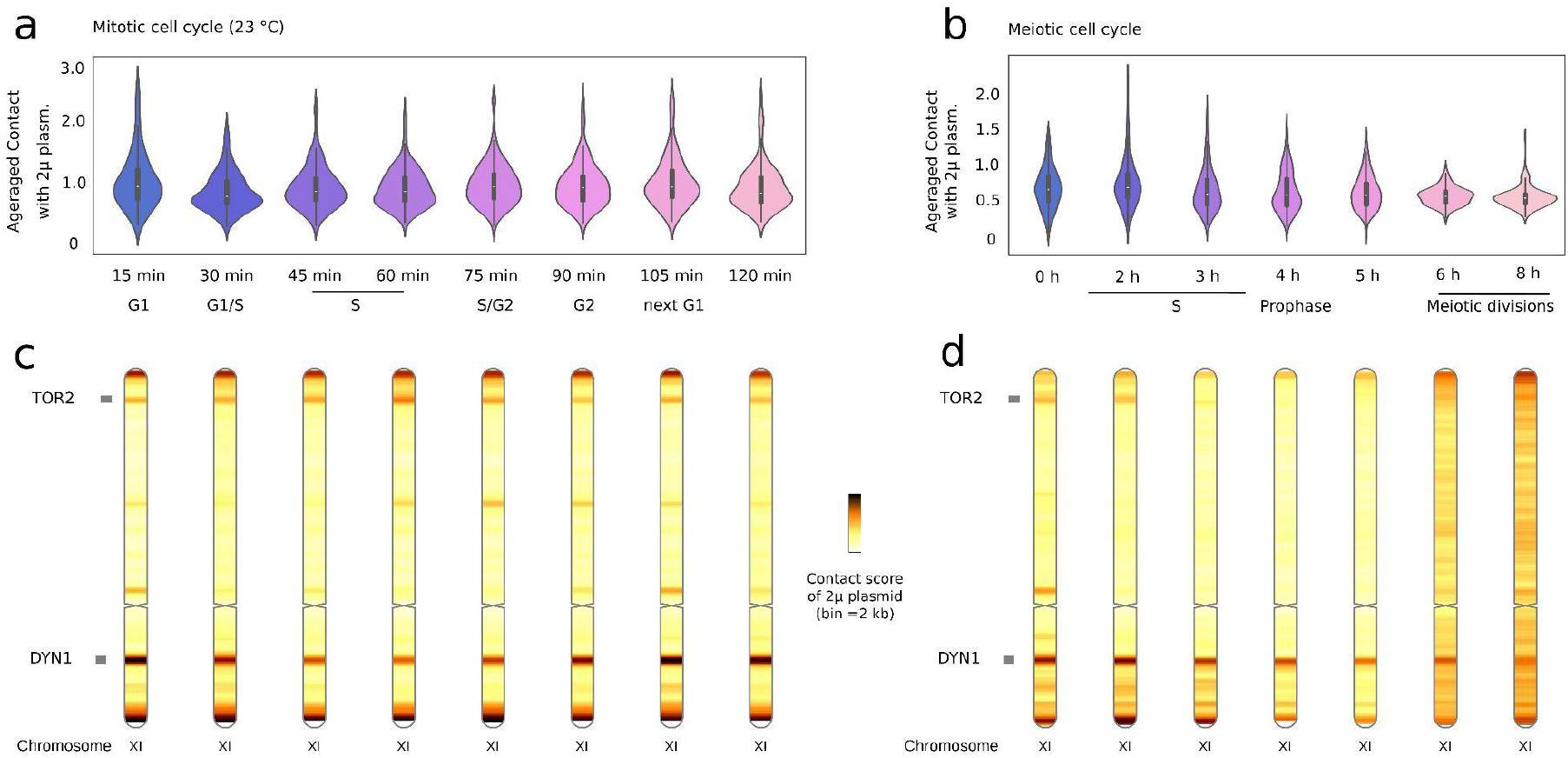
Contact of the 2*µ* plasmid during mitotic and meiotic cell cycles. **a**, Distribution of contact values of identified hotspots contacted by the 2*µ* plasmid identified in WT, log phase during the mitotic cell cycle (contact data reanalysed from [7]). **b**, Distribution of contact values of the identified hotspots contacted by the 2*µ* plasmid during the meiotic cell cycle (contact data reanalysed from [19]). **c**, Example of contact profile of 2*µ* plasmid with chromosome XI during mitotic cell cycle. **d**, Example of contact profile of 2*µ* plasmid with chromosome XI during meiotic cell cycle. Long genes (size > 7 kb) are annotated.

**Supplementary Figure 9.**
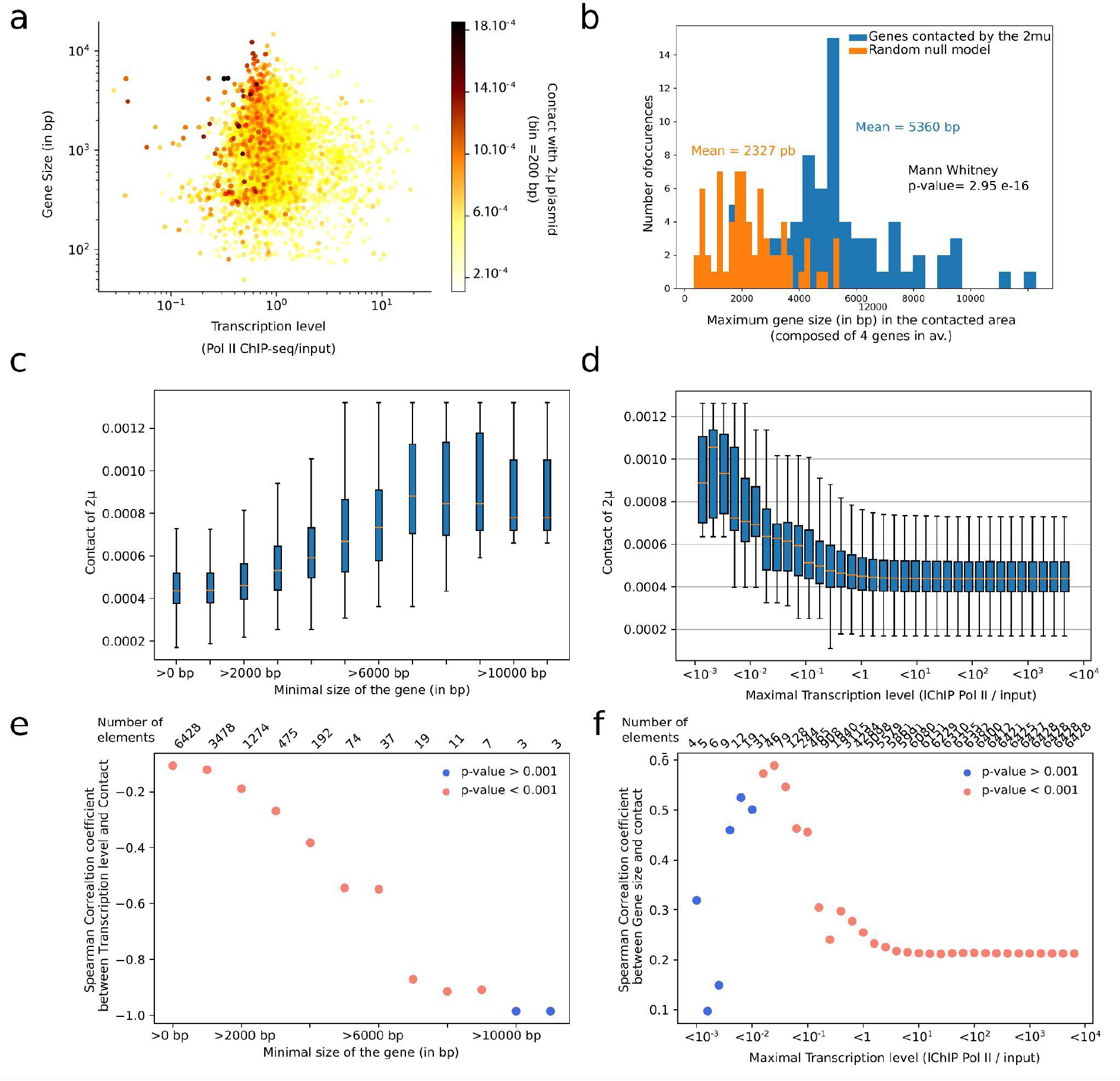
Statistical analyses on the size and transcription level of genes contacted by 2*µ* plasmid. **a**, Scatter plot for all genes of *S. cerevisiae* represented in function of their transcription level (x-axis), their size in bp (y-axis) and their level of contact with 2*µ* plasmid represented by their color (colorbar on the left). MicroC data were reanalysed from [5]. **b**, Distribution of maximum sizes of genes from loci contacted by 2*µ* plasmid and from a random group of loci with the associated statistical test. **c**, Contact level of 2*µ* plasmid in function of the minimal size of gene (in bp). **d**, Contact level of 2*µ* plasmid in function of the maximal transcription level (ChIP-seq data of Rpb3, sub-unit of PolII, [5]). **e**, Spearman correlation coefficient between transcription level and contact with 2*µ* plasmid in function of the minimal size of gene (in bp). **f**, Spearman correlation coefficient between gene size and contact with 2*µ* plasmid in function of the maximal transcription level (ChIP-seq data of Rpb3, sub-unit of PolII from [5]).

**Supplementary Figure 10.**
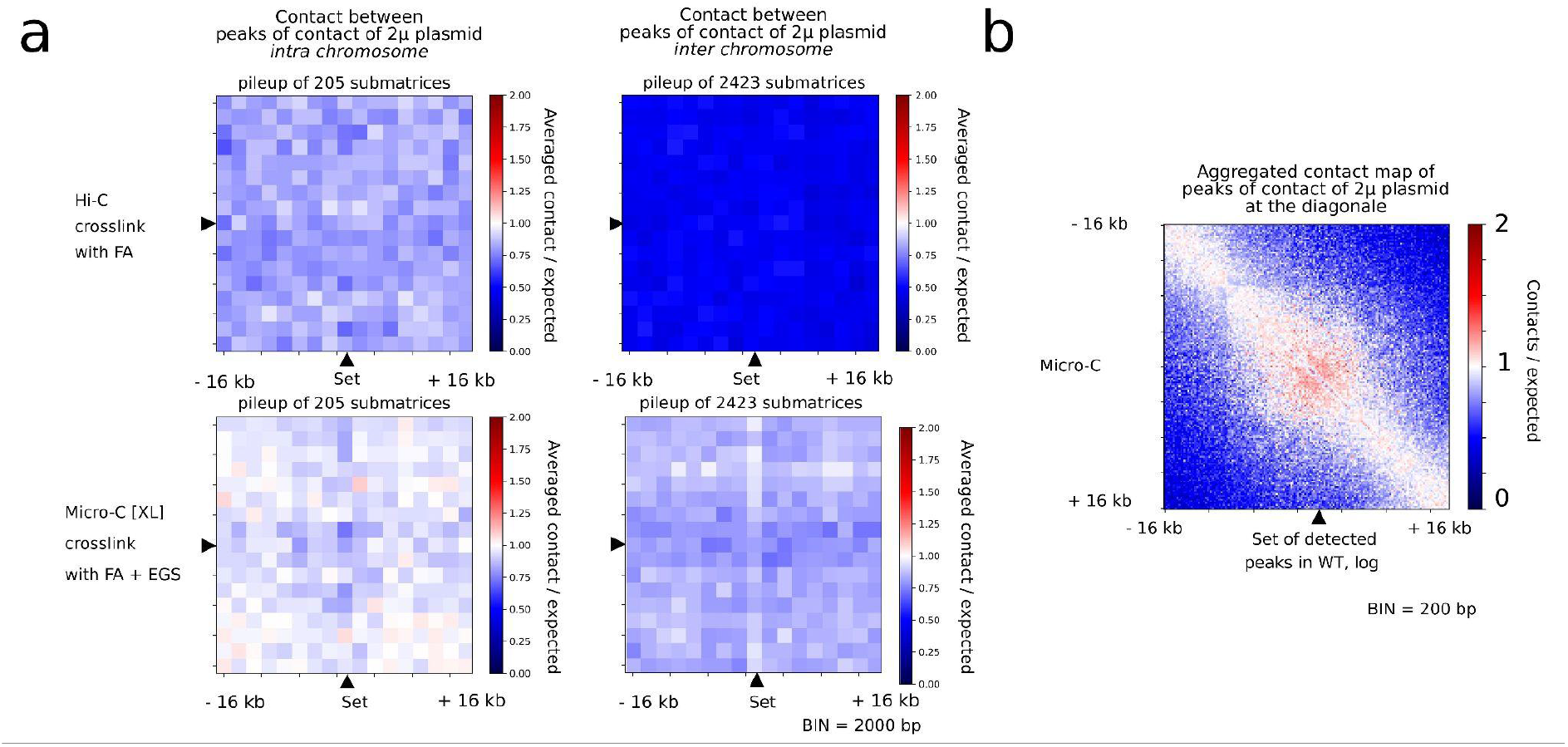
Contact behavior for the identified loci contacted by 2*µ* plasmid. **a**, Agglomerated plot between pairs of loci contacted by 2*µ* plasmid belonging to the same chromosome (left) or belonging to different chromosomes (right) for two different contact technologies: Hi-C (top) and MicroC with dual crosslink (bottom) [5]. The signal represents the ratio between the contact measured between loci contacted by 2*µ* plasmid over random pairs separated by same genomic distances [20]. **b**, Agglomerated plot at the diagonal for the identified loci contacted by 2*µ* plasmid with bins of 200 bp (MicroC data reanalysed from [5]).

**Supplementary Figure 11.**
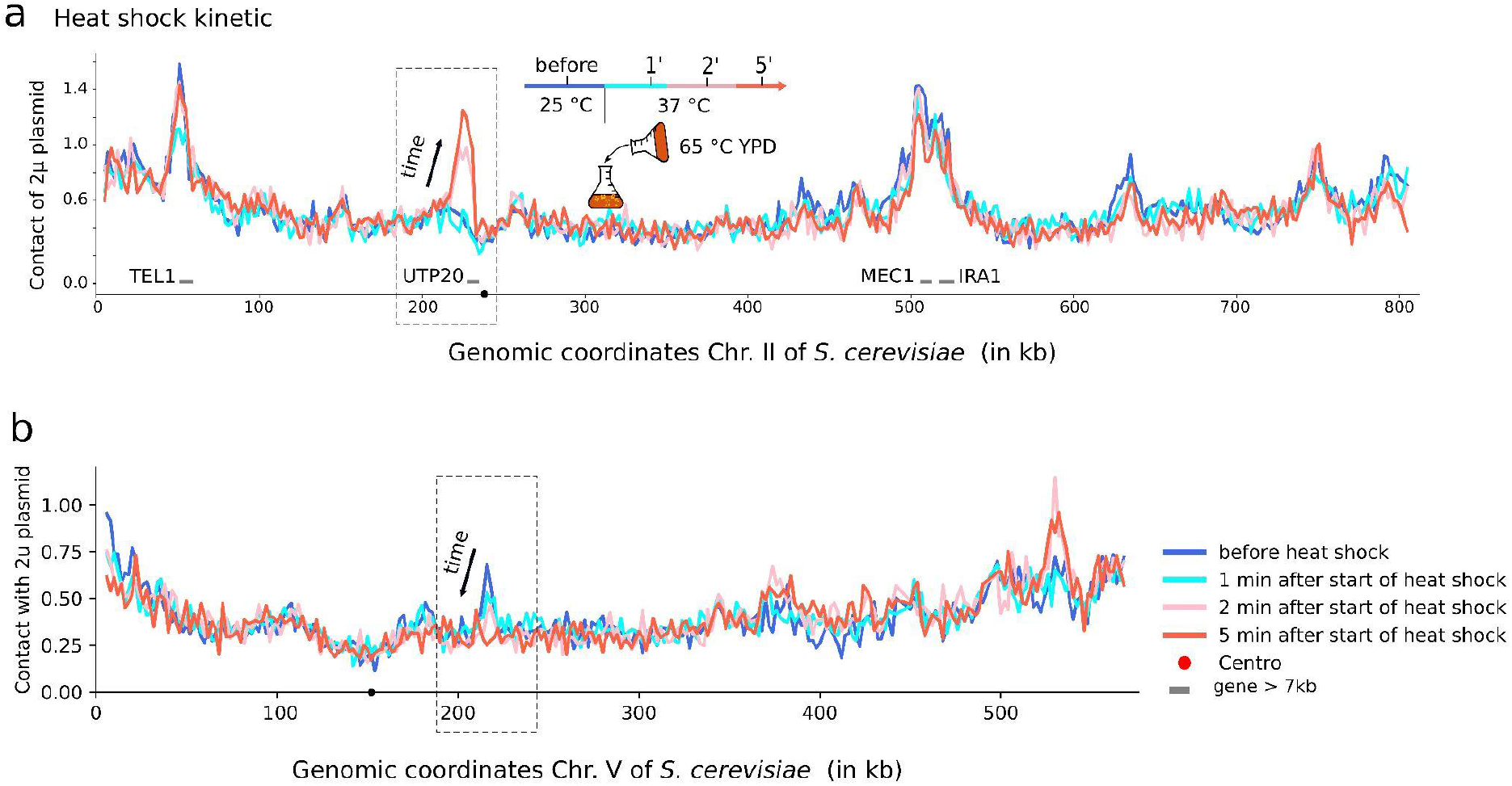
Contact signal of 2*µ* plasmid during a heat shock. **a**, Contact signal of 2*µ* plasmid along the chromosome II of *S. cerevisiae* for 4 time points: before heat shock, 1 min, 2 min and 5 min after heat shock. **b**, Contact signal of 2*µ* plasmid along the chromosome V of *S. cerevisiae* for 4 time points: before heat shock, 1 min, 2 min and 5 min after heat shock. Binning for the contact signals is 2 kb.

**Supplementary Figure 12.**
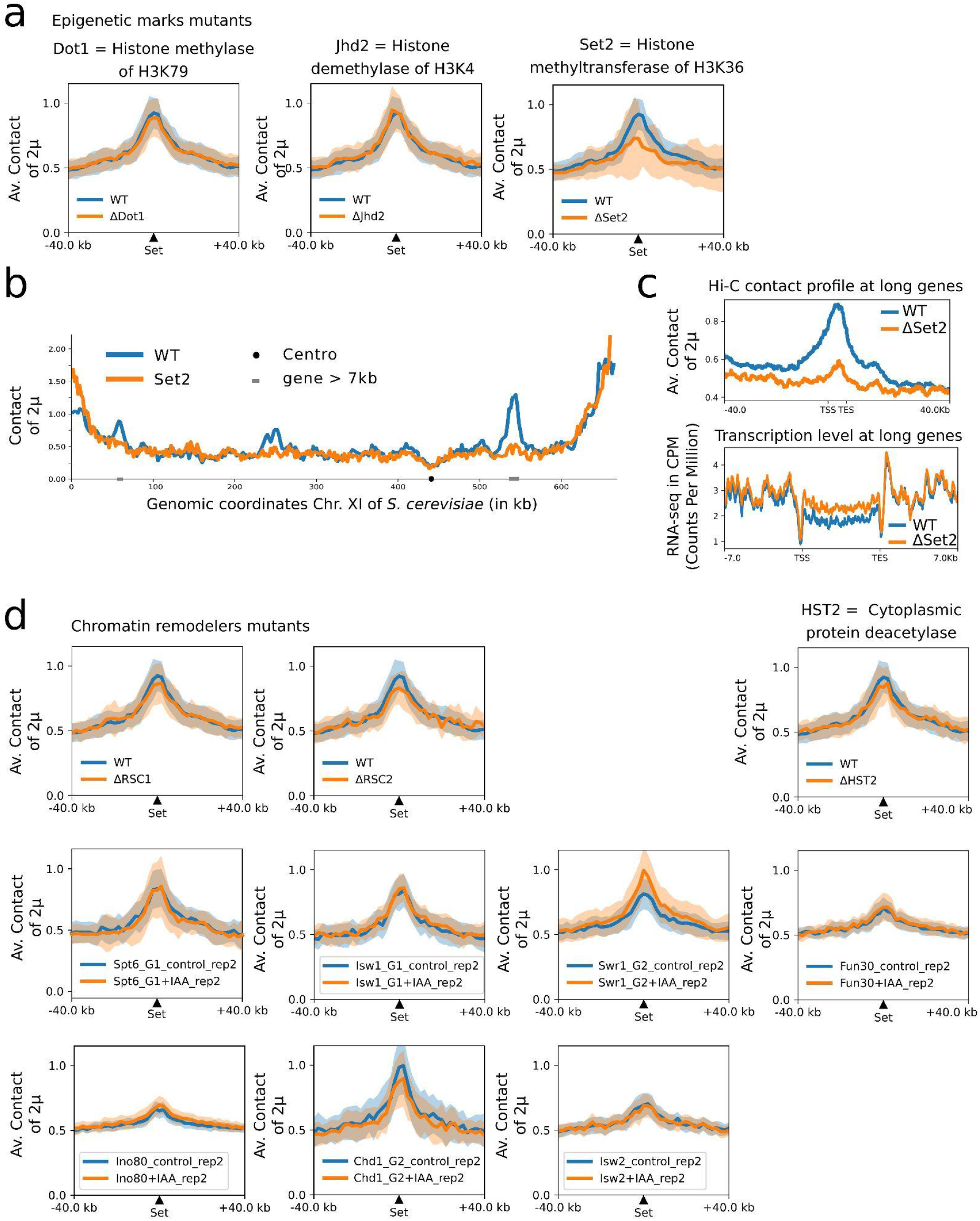
Contact signal of 2*µ* plasmid in epigenetic marks and chromatin remodelers mutants. **a**, Averaged 2*µ plasmid* contact signal over the set of identified loci contacted by 2*µ* plasmid in WT, log phase condition for ΔDot1, ΔJhd2 and ΔSet2 mutants as well as for the ΔHST2 mutant. **b**, Contact signals of 2*µ* plasmid along chromosomes VII and XII for WT and ΔSet2 mutant. **c**, Average contact signal (top) and average transcription level (below) at long genes for WT and ΔSet2 mutant. **d**, Averaged 2*µ* plasmid contact signal over the set of genomic positions identified in WT, log phase condition for ΔRSC1, ΔRSC2 as well as for 7 chromatin remodelers degradation mutants (AID system) and their corresponding control: Spt6, Isw1, Swr1, Fun30, Ino80, Chd1, Isw2 (data from [21]).

**Supplementary Figure 13.**
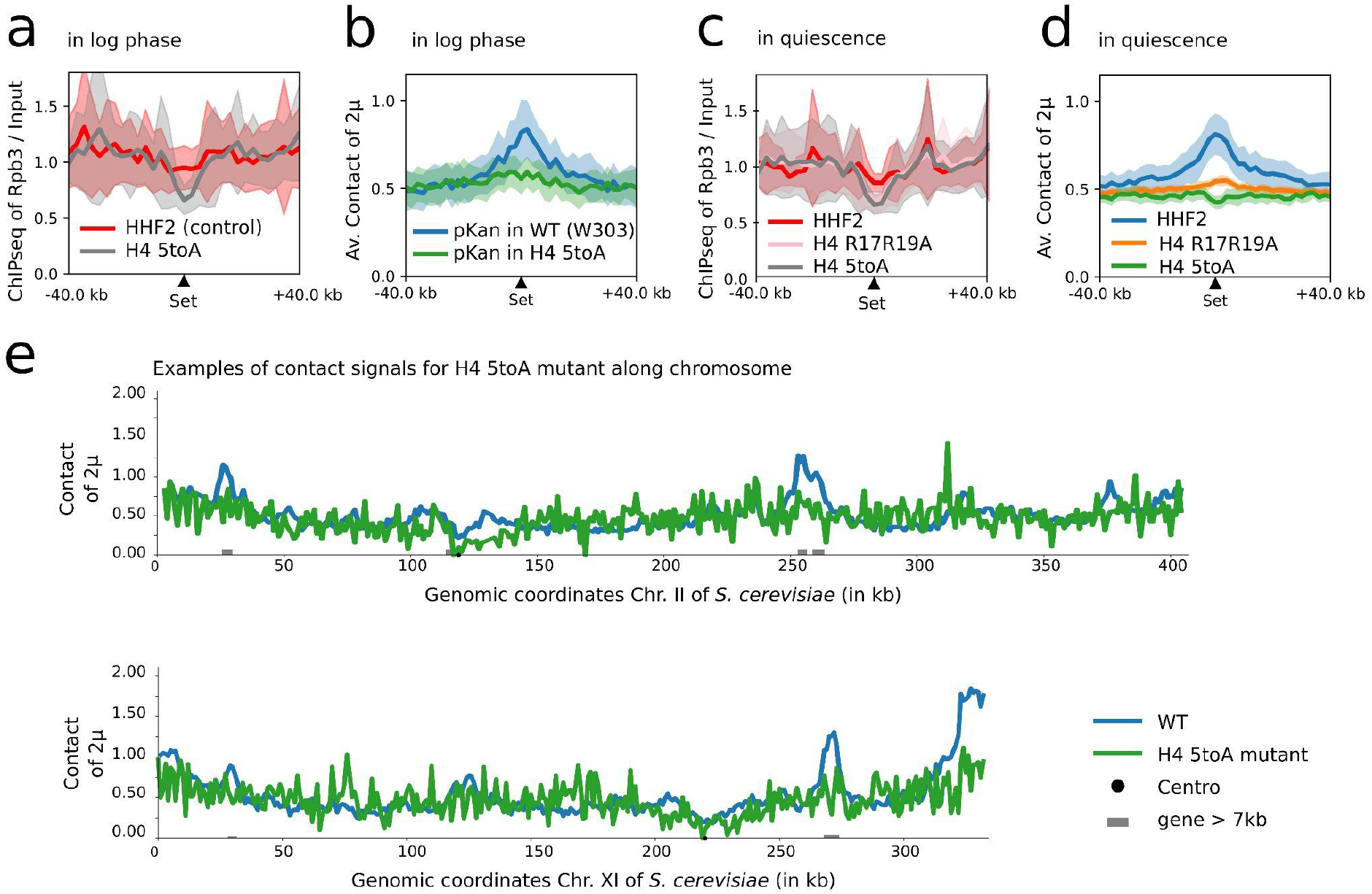
Contact signal of 2*µ* plasmid in the H4 5toA mutant. **a**, Averaged transcription signal measured by Rpb3 (PolII sub-unit) ChIP-seq [22] over the set of identified loci contacted by 2*µ* plasmid in WT, log phase condition for the control and H4 5toA mutant in log phase. **b**, Averaged 2*µ* plasmid contact signal over the set of identified loci contacted by 2*µ* plasmid in WT, log phase condition for the control and H4 5toA mutant in log phase. pKAN version of the 2*µ* plasmid was used to ensure plasmid stability. **c**, Averaged transcription signal measured by Rpb3 (PolII sub-unit) ChIP-seq [22] over the set of identified loci contacted by 2*µ* plasmid in WT, log phase condition for the control, H4 R17R19A and H4 5toA mutants in quiescence phase [22]. **d**, Averaged 2*µ* plasmid contact signal over the set of identified loci contacted by 2*µ* plasmid in WT, log phase condition for the control, H4 R17R19A and H4 5toA mutant in quiescence phase [22]. **e,** Examples of contact signals of 2*µ* plasmid (pKAN version) in WT and H4 5toA mutant.

**Supplementary Figure 14.**
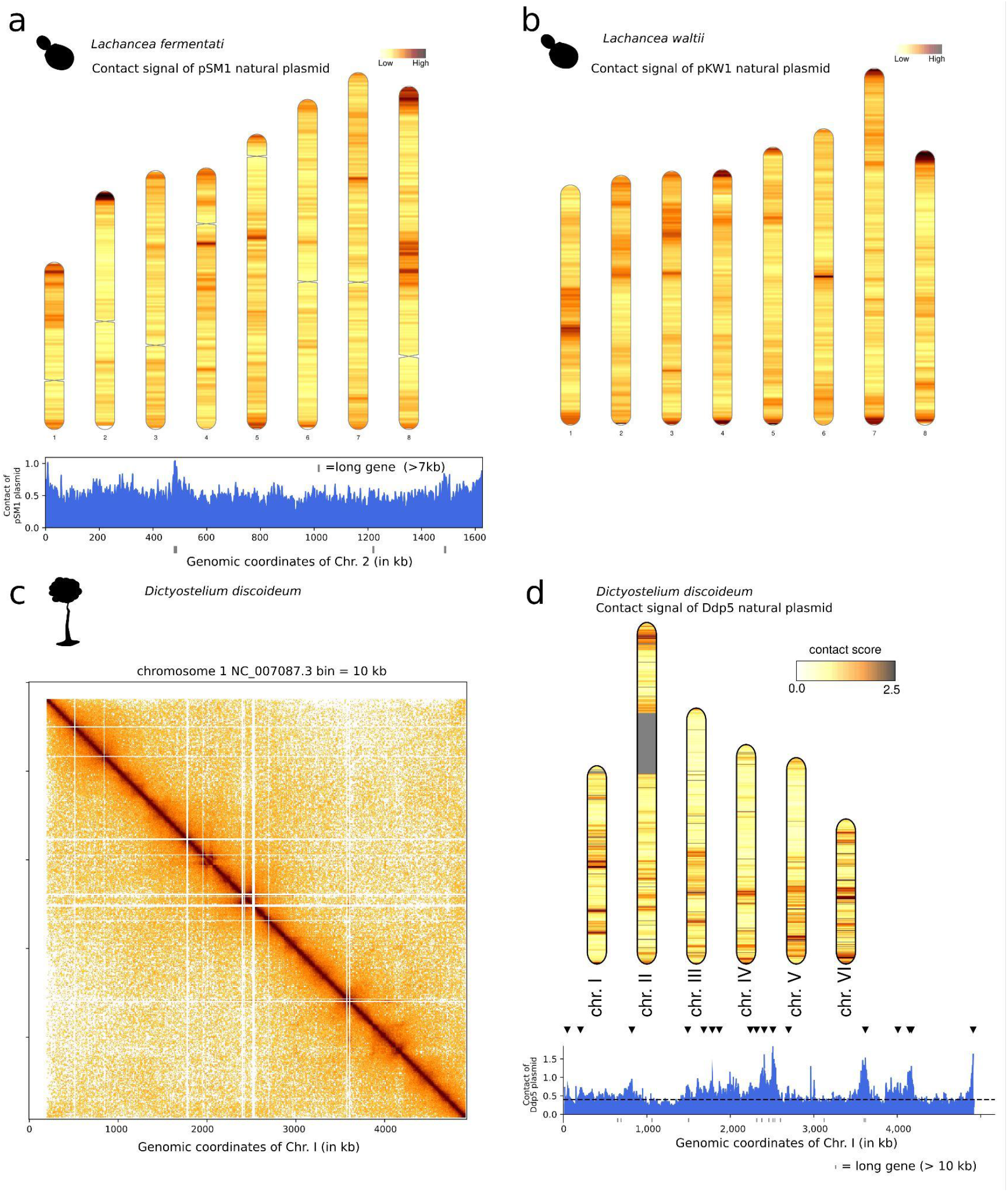
Contact signals of natural plasmids in other eukaryotes. **a**, Contact map of chromosome 1 of *Dictyostelium discoideum* amoeba binned at 10 kb resolution. **b.** Contact signal of the Ddp5 natural plasmid with the chromosomes of *Dictyostelium discoideum* and contact profile for the chromosome 1. Orange triangles indicate automatically detected peaks and long genes are annotated as grey boxes. **c**, Contact signal of the pSM1 natural plasmid with the chromosomes of *Lachancea fermentati* yeast and contact profile for chromosome 2. Long genes are annotated as grey boxes. **d**, Contact signal of the pKW1 natural plasmid with the chromosomes of *Lachancea waltii* yeast.

**Supplementary Table 1:** Contact datasets analysed in the present study. The last column indicates either the identifier for the raw reads available on the Short Read Archive server (SRA) (https://www.ncbi.nlm.nih.gov/sra) or on Gene Expression Omnibus server (GEO) https://www.ncbi.nlm.nih.gov/geo.

| **Experiment type and condition** | **Figure** | **Ref** | **Identifier** |
| --- | --- | --- | --- |
| Micro-C, WT log and quiescence phases | **Sup Fig 1,2,3** | [5] | SRR7939017<br>SRR7939018 |
| Hi-C, WT W303, G1 arrest | **Sup Fig 7** | [10] | SRR12284705 |
| MicroC, WT A364, asynchronous | **Sup Fig 7** | [7] | SRR11893084 |
| Hi-C, WT diploid heterozygous | **Sup Fig 7** | [10] | SRR12284704 |
| Hi-C, WT DSB t2 replicate2 | **Sup Fig 7** | [10] | SRR12284704 |
| Hi-C, WT, with DMSO | **Sup Fig 7** | [11] | SRR13147965 |
| Hi-C, WT, with HU | **Sup Fig 7** | [11] | SRR13147975 |
| Micro-C, Mcd1-AID (rep1) and control | **Fig1, Sup Fig 7** | [7] | SRR11893086<br>SRR11893085 |
| Hi-C, Smc2-AID and control | **Fig1, Sup Fig 7** | [12] | SRR9040342<br>SRR9040345 |
| Hi-C, Sir3-AID and control | **Sup Fig 7** | [9] | SRR12108219<br>SRR12108218 |
| Hi-C, Pds5-degron, Eco1-degron, Wapl-degron | **Sup Fig 7** | [6] | SRR10687277<br>SRR10687278 |
| Hi-C, cdc45-AID and control (alpha-factor G1) | **Sup Fig 7** | [6] | SRR10687274<br>SRR8769554 |
| Hi-C, Top2 | **Sup Fig 7** | [13] |  |
| Hi-C, mutants Chl4Δ, sgo1Δ, nocodazole treatment | **Sup Fig 7** | [14] | SRR8718857,<br>SRR8718853 |
| MicroC, Rad50 mutant and control | **Sup Fig 7** | [15] | SRR11489731<br>SRR11489732 |
| Hi-C, ΔSgs1 DSB t4, ΔSml1 DSB t4 noco, ΔMec3 DSB t4 noco | **Sup Fig 7** | [10] | SRR12284714<br>SRR12284727<br>SRR13736493 |
| Hi-C, Fkh mutant and control | **Sup Fig 7** | [16] | SRR5337954<br>SRR5337951 |
| Hi-C, Clb5-Clb6 mutant and control | **Sup Fig 7** | [17] | SRR17873477<br>SRR17873476 |
| Hi-C, Syn6.5 strain and control | **Sup Fig 7** | [23] | SRR22910279<br>SRR22910280 |
| Micro-C, mitotic cell cycle | **Sup Fig 8** | [7] | SRR11893095 to<br>SRR11893114 |
| Hi-C, meiosis (t=0h,3h,4h, 6h) | **Sup Fig 8** | [24] | SRR7126297<br>SRR7126293<br>SRR7126301<br>SRR7340033 |
| Hi-C, meiosis cell cycle, WT | **Sup Fig 8** | [19] | SRR8689946 to<br>SRR8689952 |
| Hi-C, chromatin remodelers mutants (Spt6, Isw1, Swr1, Fun30, Ino80, Chd1, Isw2 and controls) | **Sup Fig 12** | [21] | GSE158336 |

**Supplementary Table 2:** Genomic datasets (other than contact data) analyzed in the present study. The last column indicates either the identifier for the raw reads available on the Short Read Archive server (SRA) (https://www.ncbi.nlm.nih.gov/sra), the identifier of the .cool files accessible on the Gene Expression Omnibus server (GEO)https://www.ncbi.nlm.nih.gov/geo

| Experiment type and condition | Figure | Ref | Identifier |
| --- | --- | --- | --- |
| Pol II ChIP-seq | Fig 3 | [25] | SRR1916157<br>SRR1916162 |
| H3 ChIP-seq, in Log Rep1 | Fig 3 | [22] | SRR13736587 |
| RNA-seq | Fig 1,2 & Sup Fig 1 | [2]<br>[26] | SRR14693235<br>SRR7692240 |
| ATAC-seq | Fig 1 | [27] | SRR6246290 |
| ~1300 ChIP-exo covering ~800 different proteins and genomic signals | Fig 3 | [28] | GSE147927 |
| ATAC-seq | Fig1, Sup Fig 1 | [3] | SRR11235539 |
| Cse4 ChIP-seq | Sup Fig 1 | [4] | SRR10765000<br>SRR10764999 |
| Scc1 ChIP-seq | Sup Fig 1 | [29] | SRR1103930<br>SRR1103928 |
| Brn1 ChIP-seq | Sup Fig 1 | [5] | SRR7175367<br>SRR7175368 |
| RNA-seq in *Dictyostelium discoideum* (vegetative stage, rep1) | Fig 4 | [30] | SRR10133961? |

**Supplementary Table 3:** List of strains used in the present study.

| Name | genotype / specie if not *S. cerevisiae* | Origin |
| --- | --- | --- |
| BY4741 | MATa his3Δ1 leu2Δ0 met15Δ0 ura3Δ0 cir+ | Brachmann et al. 1998 |
| BY4742 | MATα his3Δ1 leu2Δ0 lys2Δ0 ura3Δ0 | Brachmann et al. 1998 |
| BY4743 | MATa/α; his3Δ1/his3Δ1; leu2Δ0/leu2Δ0; met15Δ0/MET15; LYS2/lys2Δ0; ura3Δ0/ura3Δ0 | Brachmann et al. 1998 |
| W303 | MATa {leu2-3,112 trp1-1 can1-100 ura3-1 ade2-1 his3-11,15} | R. Rhotstein |
| Y9-4 | strain from Indonesia (used for Ragi fermentation, finger millet) | Peter et al.2018 |
| BHB | strain from bakeries in Australia | Peter et al. 2018 |
| H4 5toA | MATa RAD5+ ura3-1 hht1-hhf1::Nat hht2- hhf2::Hyg trp1-1::pRS 404-HHT2-hhf2-K16A, R17A,H18A,R19A,K20A | Swygert et al 2021 |
| RSGY 712 | W303 M.mycoides linear | Chapard et al. 2023 |
| RSGY 960 | W303 [cir-] pKAN | McQuaid et al.2019 |
| RSGY 960 | BY4741 M.pneumoniae linear | Chapard et al. 2023 |
| RSGT 1056 | BY4741 jhd2::KANMX4 cir + | This study |
| RSGY 1055 | BY4741 set2::KANMX4 cir+ | This study |
| RSGY 1057 | BY4742 X RSGY 960 cir+ | This study |
| RSGY 1058 | BY4742 X RSGY 712 cir + | This study |
| RSGY 1059 | BY4741 cir- | This study |
| RSGY 1065 | Y9-4 pKAN | This study |
| RSGY 1068 | BY4741cir- pKAN-ΔREP1 | This study |
| RSGY 1069 | BY4741cir- pKAN-ΔSTB-P | This study |
| RSGY 1070 | BY4741 cir-pKAN | This study |
| RSGY 1116 | BY4741 dot1 ::KANMX4 | This study |
| RSGY 1217 | BY4741 hst2::KANMX4 | This study |
| RSGY 1218 | H4 5toA pKAN | This study |
| RSGY 1226 | BY4741 rsc1 ::KANMX4 | This study |
| RSGY 1252 | BY4741 rsc2::KANMX4 | This study |
| RSGY 1255 | BY4741 cir- pARS | This study |
|  | *Lachancea waltii* | Gilles Fisher |
|  | *Lachancea fermentati* | Gilles Fisher |
| AX4 | *Dictyostelium discoideum* | DBS0237907 |
| yAT4593 | MATalpha rap1::RAP1-GFP(ADE2) (W303 background) | Angela Taddei |
| yAT4595 | MATa rap1::RAP1-GFP(ADE2) XVI-mycoides-fusion (W303 background) | Angela Taddei |

**Supplementary Table 4:** List of plasmids used in this study.

| Name | Relevant genetic features | Origin |
| --- | --- | --- |
| pKAN ΔREP1 | pKAN ::ΔREP1 | McQuaid et al.2019 |
| pKAN ΔSTB | pKAN ::ΔSTB-P | McQuaid et al.2019 |
| pKAN-STB-P | REP1 REP2 STB-P FLP1:KANMX4:FLP1<br>IR1 IR2 RAF1 | McQuaid et al.2019 |
| pARS | pRS413 ::CEN4 | This study |
| pRS413 | CEN4 ARS HIS3 | Sikorksi et al. 1989 |

## Notes

### Competing Interest Statement

The authors have declared no competing interest.

